# Permissive central tolerance plus defective peripheral checkpoints licence pathogenic memory B cells in CASPR2-antibody encephalitis

**DOI:** 10.1101/2025.01.14.631703

**Authors:** Bo Sun, Dominique Fernandes, Anne-Kathrin Kienzler, Sofija Paneva, Ruby Harrison, Sudarshini Ramanathan, Anna L Harrison, Mateusz Makuch, Miriam L Fichtner, Robert F. Donat, Deniz Akdeniz, Halwan Bayuangga, Min Gyu Im, Robyn Williams, Ana Vasconcelos, Selina Thomsen, Andrew Fower, Ruyue Sun, Hannah Fox, Victor Mgbachi, Alexander Davies, Mandy Tseng, Adam Handel, Mark Kelly, Meng Zhao, James Bancroft, Rachael Bashford-Rogers, John V Pluvinage, Ravi Dandekar, Bonny D. Alvarenga, Lynn Dustin, Simon Rinaldi, Ray Owens, Daniel Anthony, David L Bennett, Patrick Waters, Simon J. Davis, Michael R Wilson, Kevin C O’Connor, John Soltys, Ana Luisa Carvalho, Sarosh R Irani

## Abstract

Autoimmunity affects 10% of the population. Within this umbrella, autoantibody-mediated diseases targeting one autoantigen provide a unique opportunity to comprehensively understand the developmental pathway of disease-causing B cells and autoantibodies. While such autoreactivities are believed to be generated during germinal centre reactions, the roles of earlier immune checkpoints in autoantigen-specific B cell tolerance are poorly understood. We address this concept in patients with CASPR2-autoantibody encephalitis and healthy controls. In both groups, comparable and high (∼0.5%) frequencies of unmutated CASPR2-reactive naïve B cells were identified. By contrast, CASPR2-reactive memory B cells were exclusive to patients, and their B cell receptors demonstrated affinity-enhancing somatic mutations with heterogenous binding kinetics. These effector molecules possessed epitope-dependent pathogenic effects *in vitro* neuronal cultures and *in vivo.* The unmutated common ancestors of these memory B cells showed a distinctive balance between strong CASPR2 reactivity and very limited binding across the remaining human proteome. Our results are the first to propose mechanisms underlying autoantigen-specific tolerance in humans. We identify permissive central tolerance, defective peripheral tolerance and heterogenous autoantibody binding properties as sequential pathogenic steps which licence CASPR2-directed pathology. By leveraging the basic immunobiology, we rationally direct tolerance-restoring approaches in CASPR2-antibody diseases. This paradigm is applicable across autoimmune conditions.

## Introduction

Across autoimmunity, few diseases are proven to be mediated by autoantibodies which target a single antigen.(1) A comprehensive understanding of the developmental autoantigen-reactive B cell lineage, and their escape from immune checkpoints, is likely sufficient to fully explain the pathway to disease causation.(2, 3) As such, prototypical autoantibody-mediated conditions provide unique biological and translational opportunities.

The recent discovery of several causative autoantibodies which target cell surface neuronal proteins has revolutionized the diagnosis of multiple neurological conditions, most notably forms of autoimmune encephalitis (AE).(4) One such protein is contactin-associated protein-like 2 (CASPR2). Autoantibodies against the extracellular domain of CASPR2 associate with a common form of AE (CASPR2-Ab-E) which presents with memory loss, behavioural disturbances, seizures, cerebellar dysfunction and neuropathic pain, consistent with the expression of CASPR2 in both central and peripheral nervous systems.(5-8) The direct pathogenicity of CASPR2-antibodies is supported by passive transfer of polyclonal patient serum IgG to rodents, which reproduces core clinical features observed in CASPR2-Ab-E patients.(9, 10) Despite some improvements with immunotherapies, nearly all patients with CASPR2-Ab-E remain disabled by multiple residual neuropsychiatric deficits or persistent neuropathic pain.(5-8, 11) Further, around 40% of patients relapse despite immunotherapies.(6, 7) Current treatment options remain limited to broad-acting immunotherapies including corticosteroids, rituximab, and intravenous immunoglobulins.(5, 7, 11-13) Precise immunotherapeutic paradigms for AE are needed to prevent the accumulation of irreversible neurologic dysfunction, and to mitigate adverse effects commonly encountered with available immunotherapies.

To this end, a better understanding is required of the underlying immunological mechanisms driving CASPR2-Ab-E. The origins and inadvertent escape of autoreactive B cells form the fundamental pathway to pathogenic autoantibody production. In this process, key immune checkpoints need to be traversed by autoreactive B cells.(14) These include a ‘central tolerance’ checkpoint, governing bone marrow exit and entry to the circulating naïve B cell (NBC) compartment.(15) Thereafter, several peripheral checkpoints likely oversee entry into the later NBC and memory B cell (MBC) and plasma cell repertoires.(15-17) Examples of how autoantigen-specific B cells are tolerised in these processes are limited, with no mechanistic exploration in neurological disorders to date.

Nevertheless, a few clues have emerged. Paradigms from non-neurological and neurological autoantibody-mediated diseases suggest that the autoantigen specificity of MBC-derived B cell receptors (BCRs) is lost when BCR somatic hypermutations are reverted to their unmutated common germline ancestors (UCAs).(18-20) Although not a universal finding,(21) this observation strongly implicates germinal centres as key sites which generate higher-affinity, pathogenic autoantigen-reactive BCRs. In contrast, more recent evidence in autoantibody-mediated neurological diseases suggest a more prominent role for autoantigen-specific NBCs. NBC BCRs recognise their cognate autoantigen aquaporin-4 in patients with neuromyelitis optica spectrum disorder,(22) and autoantigen-specific unmutated BCRs with pathogenic potential have been detected in the cerebrospinal fluid of patients with N-methyl-D-aspartate receptor antibody encephalitis.(23) These findings led us to hypothesise that early loss of B cell tolerance may represent an underappreciated phenomenon in autoantibody-mediated neurological conditions. Further, as CASPR2-antibodies have been reported in sera from healthy individuals and disease controls,(24) we reasoned CASPR2-Ab-E represents an elegant paradigm to evaluate how B cells traverse immune checkpoints in both health and disease.

In this study, we isolated 37 CASPR2-reactive BCRs from both NBCs and MBCs across CASPR2-Ab-E patients and healthy controls (HCs), and compared their frequencies, biophysical characteristics, and functional properties. Our findings are the first to describe the precise balance of autoantigen reactivities in human BCRs which facilitate escape from key checkpoints. Hence, we inform the earliest fundamental events in the development of causative pathogenic CASPR2-reactive BCRs and establish mechanisms underlying dysregulated B cell tolerance as a plausible rationale for novel tolerance-restoring therapeutics with relevance across multiple autoantigen-directed immune conditions.(25)

## Results

### CASPR2-reactive central and peripheral tolerance defects in health and disease

To investigate the integrity of pre- and post-germinal centre tolerance to CASPR2-reactive BCRs in both health and disease, either 10^5^ NBCs (CD19^+^IgD^+^CD27^-^) or MBCs (CD19^+^IgD^-^ CD27^+^) were isolated from six CASPR2-Ab-E patients and six HCs, and bulk cultured under conditions that promote IgM and IgG secretion (Fig.1A, Table S1). Culture supernatants were screened for human CASPR2-reactivity using a live cell-based assay, employing HEK293T cells surface expressing full-length human CASPR2. Bulk NBC supernatants from CASPR2-Ab-E patients and all HCs contained IgMs which bound the extracellular domain of CASPR2 (Fig.1B). In contrast, only MBC supernatants from patients contained CASPR2-IgG. Supernatants displayed no reactivity towards two other central nervous system autoantigens, LGI1 and aquaporin-4 (Fig. S1A).

**Figure 1:**
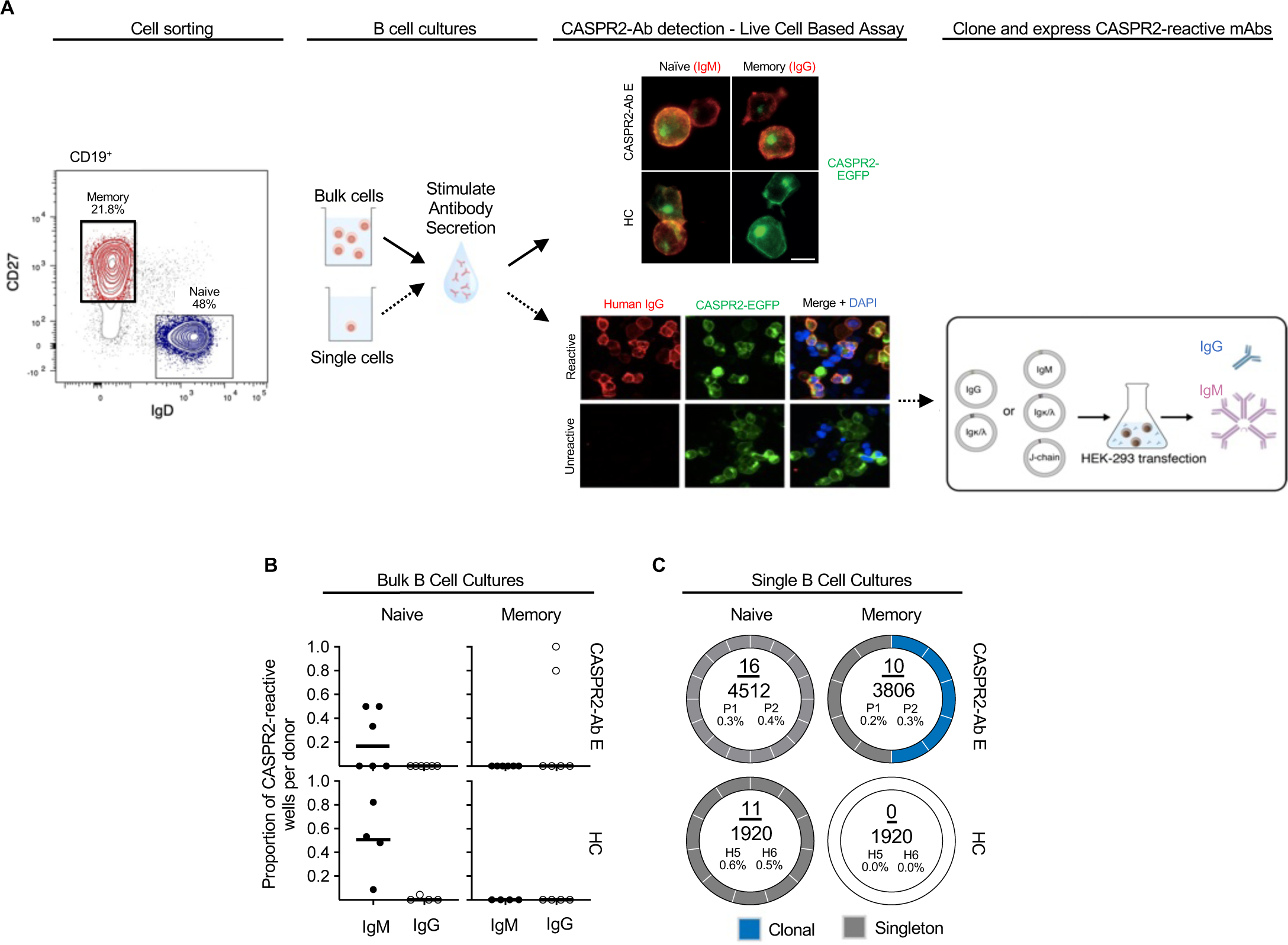
Central and peripheral immune tolerance in CASPR2-antibody encephalitis and healthy controls. (A) Left: Representative flow cytometry cell gating strategy to isolate memory (red) and naïve (blue) B cells for bulk (top) and single (bottom, dotted line) B cell cultures. Representative fluorescence microscopy images of culture supernatant detection of secreted CASPR2-reactive IgG or IgM, using a live cell-based assay with CASPR2-EGFP expressing HEK293T cells. DAPI = nuclear stain; scale bar = 10μm. Right: mRNA was extracted from CASPR2-reactive single B cell cultures to amplify and clone paired heavy and light chain BCR sequences. These were expressed in HEK293S cells to secrete CASPR2-reactive IgG (blue) or IgMs (pink). (B) The proportion of bulk B cell culture wells containing CASPR2-reactive IgM or IgG from patients (n=6) and healthy controls (HC; n=5). Black bar depicts median value. (C) Donut plots visualizing the frequency and clonality of CASPR2-reactive BCRs in single cell cultures. Numerator= total number of CASPR2-reactive BCRs, denominator = total number of cells screened. The absolute percentage of CASPR2-reactive BCRs is then shown for two patients (P1 and P2) and two HCs (H5 and H6).

To enumerate this CASPR2-reactivity at the individual B cell level and, in parallel, isolate corresponding cognate-paired heavy and light chain BCR sequences, we performed single B cell cultures.(26) 12,158 single NBCs or MBCs from the blood of two, untreated CASPR2-Ab-E patients and two HCs were sorted, (Fig.S1B) cultured, and supernatants screened for CASPR2 reactivities (Fig.1A, Table.S1). Aligned with the bulk culture findings, CASPR2-IgMs were detected in both patient and healthy donor NBC supernatants at similar frequencies (16/4512 (0.4%) vs 11/1920 (0.6%); p=0.21), while CASPR2-IgG was detected exclusively in patient MBCs (10/3806 (0.26%) versus 0/1920; p=0.037, Fisher’s exact test; Fig.1C). After sequencing, two clonal populations were observed within the MBCs (Fig.1C; Table.S2). Next, CASPR2 monoclonal antibodies (mAbs) were generated from all culture wells with CASPR2-reactive supernatants by cloning cognate-paired heavy and light chain BCR sequences into expression plasmids. As expected, all mAbs bound the extracellular domain of human CASPR2 (Fig.1A).

These concordant results across bulk and single B cell cultures suggested that ∼0.5% of NBCs harbour CASPR2-reactive BCRs in both health and disease. However, from memory compartments, CASPR2-reactive BCRs were exclusively detected in patients, and included clonal expansions.

### Origins of CASPR2 autoreactivity

Further BCR sequence analyses revealed that all NBC heavy and light chain variable regions, in both patients and HCs, contained no or very few mutations when compared to reported ancestral BCR gene segments (median 0 nucleotides, range 0-2; Fig.2A, Table.S2). In distinction, memory BCRs were all mutated, often highly so (median 22 nucleotides, range 5-38, for heavy chains; median 17, range 4 – 29 for light chains).

**Figure 2:**
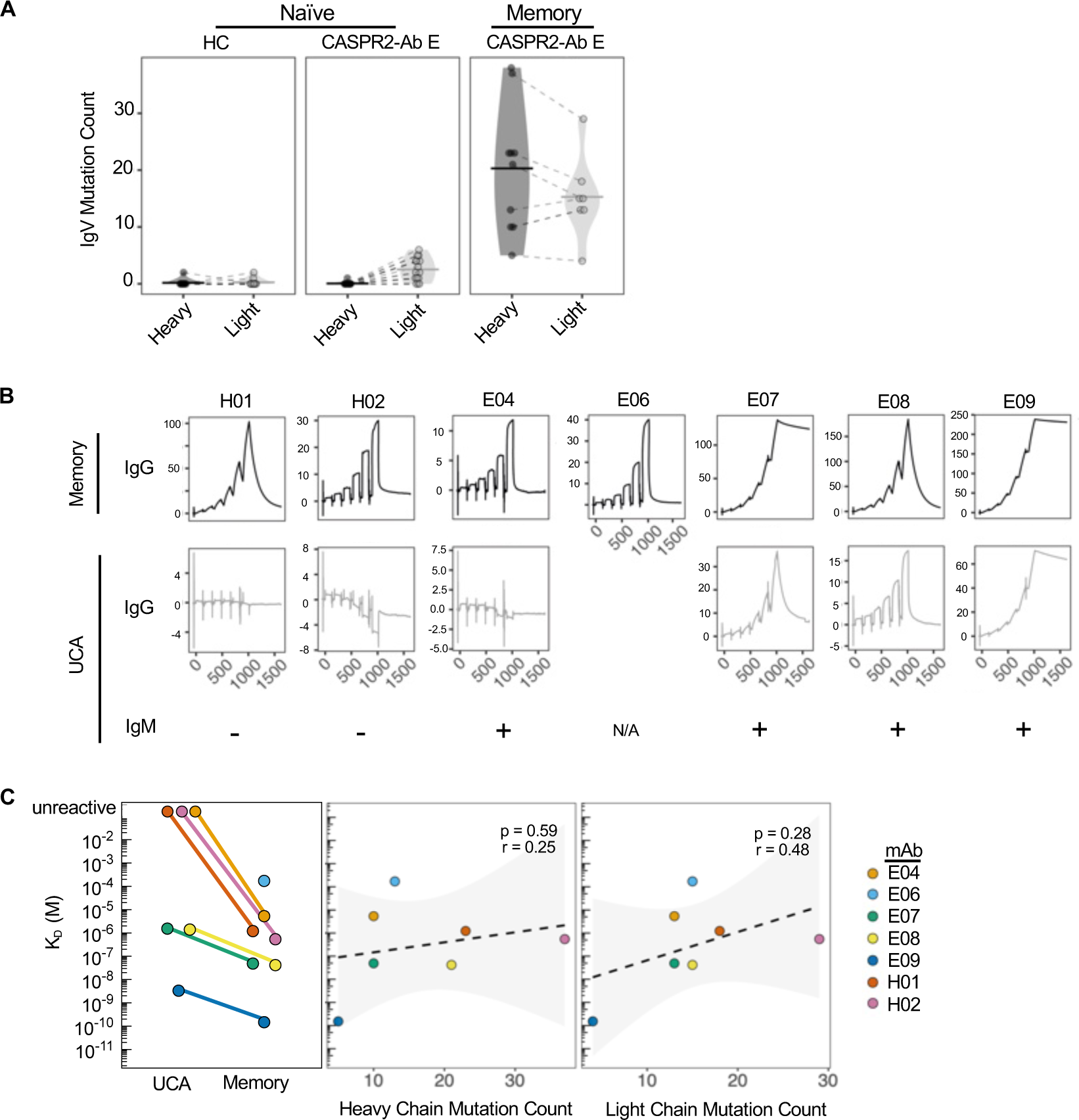
CASPR2 autoreactivity is enhanced by somatic hypermutation. (A) Immunoglobulin heavy and light chain variable region mutation counts across B cell subsets in both patients (CASPR2-Ab-E) and healthy controls (HC). (B) Raw SPR traces representing the soluble extracellular domain of human CASPR2 binding to immobilized CASPR2 memory mAbs (upper row) and their corresponding UCAs (middle row). E06 UCA mAb did not express. UCA binding as an IgM was tested via a live cell-based assay with “+” indicating CASPR2-reactivity (bottom row). (C) K_D_ (M) quantification of mAbs via surface plasmon resonance (left). Non-significant K_D_ Pearson’s correlations with heavy (middle) and light (right) chain mutation count.

To explore how somatic hypermutation affected CASPR2 reactivity, memory BCRs were reverted to their corresponding UCAs. mAb affinity was determined by binding the extracellular domain of human CASPR2 to protein A-immobilised mAbs, using surface plasmon resonance (SPR). Memory mAb affinities varied by 10^6^-fold and included some in the ‘high affinity’ picomolar range (K_D_: 171 µM to 152 pM); their on and off rates were similarly heterogeneous (Fig.2B and Table.S3). When expressed as UCAs, two mAbs (H01 and H02) lost detectable binding to CASPR2, both in IgG and pentameric IgM formats (Fig.2B). In addition, one mAb (E04) lost reactivity as an IgG but binding was detectable as an IgM. Overall, by comparison to memory counterparts, all UCAs showed a mean worsening of 29-fold in K_D_ (range 1.4 µM to 3.33 nM; Fig.2C, left panel). K_D_ did not correlate with either heavy or light chain absolute mutation counts (Fig.2C, middle and right panels).

Taken together, and consistent with the *ex vivo* isolation of naïve CASPR2-reactive BCRs, unmutated germline ancestors derived from MBCs usually retained detectable binding to CASPR2. However, somatic hypermutation conferred both improved CASPR2 reactivity and affinities, indicating the importance of mutations in generating the highest affinity CASPR2 binders.

### MBC-derived mAbs bind discrete conformationally-dependent domains on native CASPR2

Having established binding kinetics, we determined other core antigenic properties which affect binding of potentially pathogenic memory mAbs. First, to understand their preferred conformation for CASPR2, mAbs were used to immunoprecipitate linearized 49-mer peptides tiled across the full length of CASPR2 (Fig.3A).(27, 28) None showed greater binding than isotype control mAbs. Further, from Western blotting, only one of seven bound to denatured and reduced CASPR2 at high mAb concentrations (Fig.S2A). Hence, these mAbs did not bind linearised or short regions of CASPR2. Rather, and consistent with their preference for native CASPR2, binding was observed upon their application to lightly fixed and unfixed neuronal substrates which represent the most relevant anatomical localisations of symptoms in CASPR2-Ab-E patients: hippocampus, cerebellum, dorsal root ganglia and peripheral sensory neurons (Fig.3B and Fig.S2B-C). Indeed, 4/7 memory mAbs bound mouse dorsal root ganglia and brain sections, and 3/7 bound the surface of live rat hippocampal neurons and live human iPSC-derived sensory neurons. Binding was abolished upon immunostaining CASPR2^-/-^ (knockout) brain sections, confirming exclusive CASPR2 specificity on brain tissue. No NBC-derived CASPR2-reactive mAbs bound these substrates (Fig.S2C).

**Figure 3:**
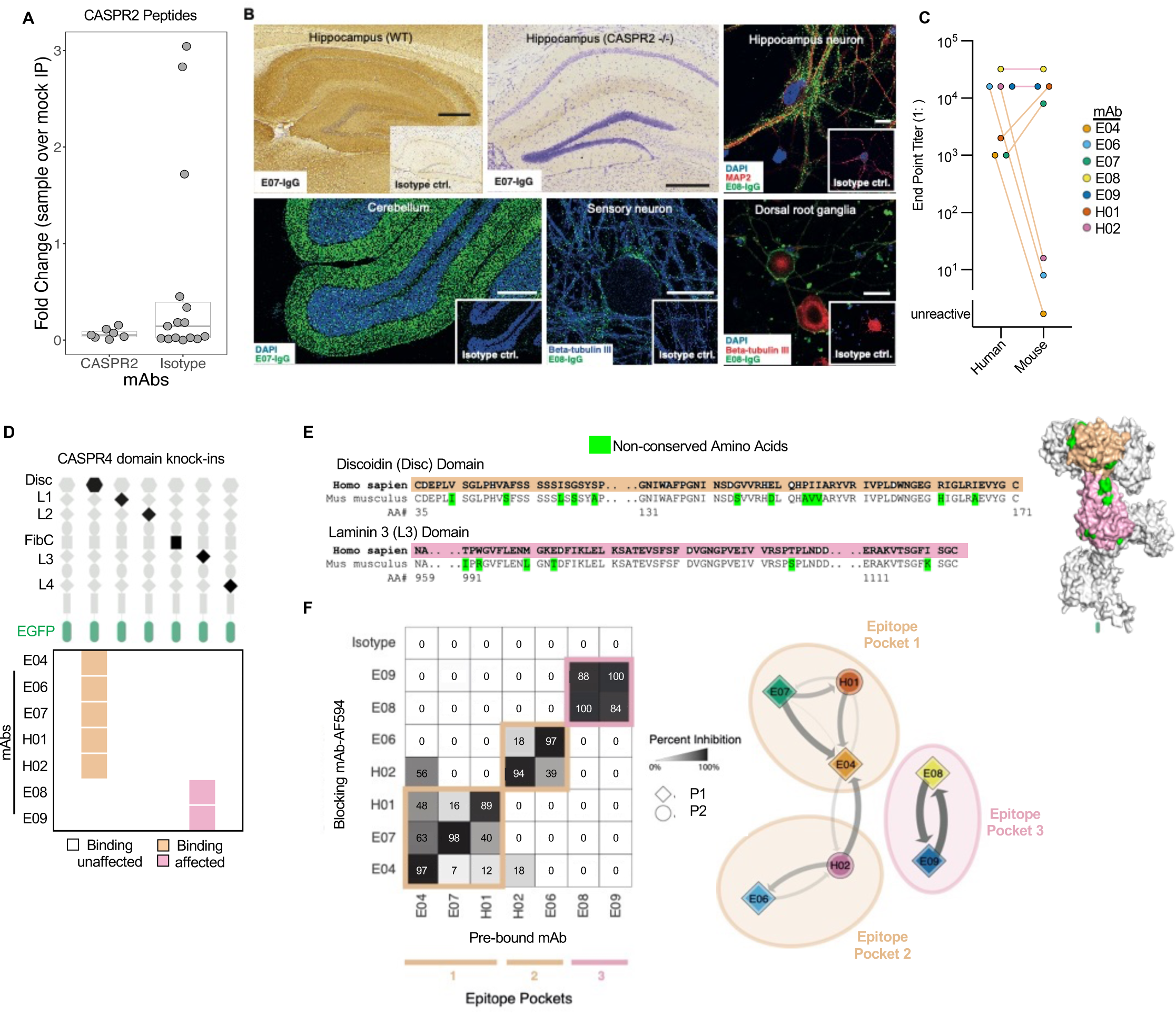
CASPR2 mAbs bind distinct conformational epitopes in native tissues. (A) No differences in number of CASPR2 peptides immunoprecipitated by peptide phage display versus isotype control mAb. (B) Representative immunohistochemistry staining (all inlays = isotype control mAb) of CASPR2 mAbs on fixed murine hippocampal brain tissue with hippocampus visualized (upper left; mAb=E07) in wild type (WT) and CASPR2 knockout (-/-; upper middle) tissue. Representative immunofluorescent staining using E08 on live hippocampal neurons (upper right) fixed rat cerebellum (lower left; DAPI = nuclear counterstain), live human iPSC-derived sensory neurons (lower middle; mAb=E08), and live mouse dorsal root ganglia (lower right; mAb=E08). Costaining markers to identify cell types include microtubule associated protein 2 (MAP2) and beta-tubulin-III. Brain tissue scale bar = 500 µm, all others = 10 µm. (C) End-point titrations (1: dilutions) of binding across human and mouse CASPR2 live cell-based assays. mAbs are colored as in subpanel F. (D) Cartoon representation of CASPR4 single domains knocked-into full length human CASPR2-EGFP (top). Heatmap depicts CASPR4 knock-in domains that abrogated mAb binding (bottom). (E) Discoidin (Disc) and Laminin3 (L3) domain amino acid sequences from human and mouse, showing non-conserved amino acids in green. The tan and pink highlight Disc and L3 domains throughout the figure, respectively. Predicted CASPR2 protein structure (right, alphafold ID 5Y4M) showing non-conserved amino acids (green), Disc and L3 domains. (F) Binding competition map (left) demonstrating displacement of a prebound mAb (x-axis) by a competing mAb pre-conjugated with Alexafluor 594 (y-axis). Percentage inhibition is defined as percentage reduction of fluorescence intensity of the respective mAb compared to isotype control and represented by a forced directed network of epitope binning (right). Arrowhead indicates direction of binding competition and line thickness and intensity denote percentage inhibition. Shapes denote patient sample.

As tissue-binding mAbs were not consistently those with higher affinities, we explored if species differences in CASPR2 structure influenced mAb binding. A direct comparison of binding to the extracellular domains of surface expressed human versus mouse CASPR2 revealed distinctive patterns: three mAbs with dramatically reduced binding to mouse CASPR2, two with 10-fold increases, and two with no differences in end-point dilutions (Fig.3C). We hypothesised these differences related to epitopes. To investigate this, we first identified individual domains preferentially targeted by the mAbs using membrane-expressing fusion constructs engineered with six single-domain substitutions, by ‘knocking-in’ the structurally similar CASPR4 domain to closely preserve the native conformations of the remaining CASPR2 domains (Fig.3D). These constructs resolved binding of 5/7 mAbs to the discoidin domain, and 2/7 to the laminin G-like 3 domains. As the latter two mAbs were also the two which displayed similar binding to human and mouse CASPR2, we predicted these bound inter-species conserved regions of the laminin G-like 3 domain (Fig.3E). In contrast, the former mAbs likely bound non-conserved amino acids on the discoidin domain (Fig.3E, Fig.S2D).(29) Indeed, direct cross-competition of pairs of mAbs against human CASPR2 using fluorophore-conjugated and unconjugated mAbs refined binding to three epitope pockets: two within the discoidin domain and one within the laminin G-like 3 domain (Fig.3F). These epitopes corresponded closely to the cross-species binding differences, with consistencies across the two patients.

Hence, patient-derived mAbs with diverse kinetics show preferential binding to the extracellular domain of natively-expressed CASPR2, and principally target three regions within two domains.

### MBC-derived mAbs harbour diverse pathogenic potentials

Next, to understand whether these varied binding characteristics translated to functional heterogeneity, the individual relative pathogenic effects of mAbs were directly studied with a focus on published works using polyclonal human CASPR2-antibody sera: namely CASPR2 internalization,(30, 31) modulation of AMPAR expression and function,(32) and altered rodent behaviors.(9)

First, CASPR2 mAbs were labelled with pHrodo, a dye that fluoresces upon entry to acidic endophagosomes. After four hours of mAb incubation with HEK293 cells expressing the human CASPR2-intracellular EGFP fusion construct, all mAbs showed varied magnitudes of pHrodo signal that consistently co-localised with EGFP, representing consistent yet differential co-internalisation of the autoantibody-autoantigen complex (Fig.4A). No internalization was observed with an isotype control and all mAbs showed reduced internalization after pharmacologic inhibition of dynamin. The magnitude of internalization did not correlate with K_D_, K_on_, or K_off_, but more closely associated with the three established epitope pockets (Fig.3F), with mAbs directed against pocket 3 showing greatest internalisation (Fig.4B).

**Figure 4.**
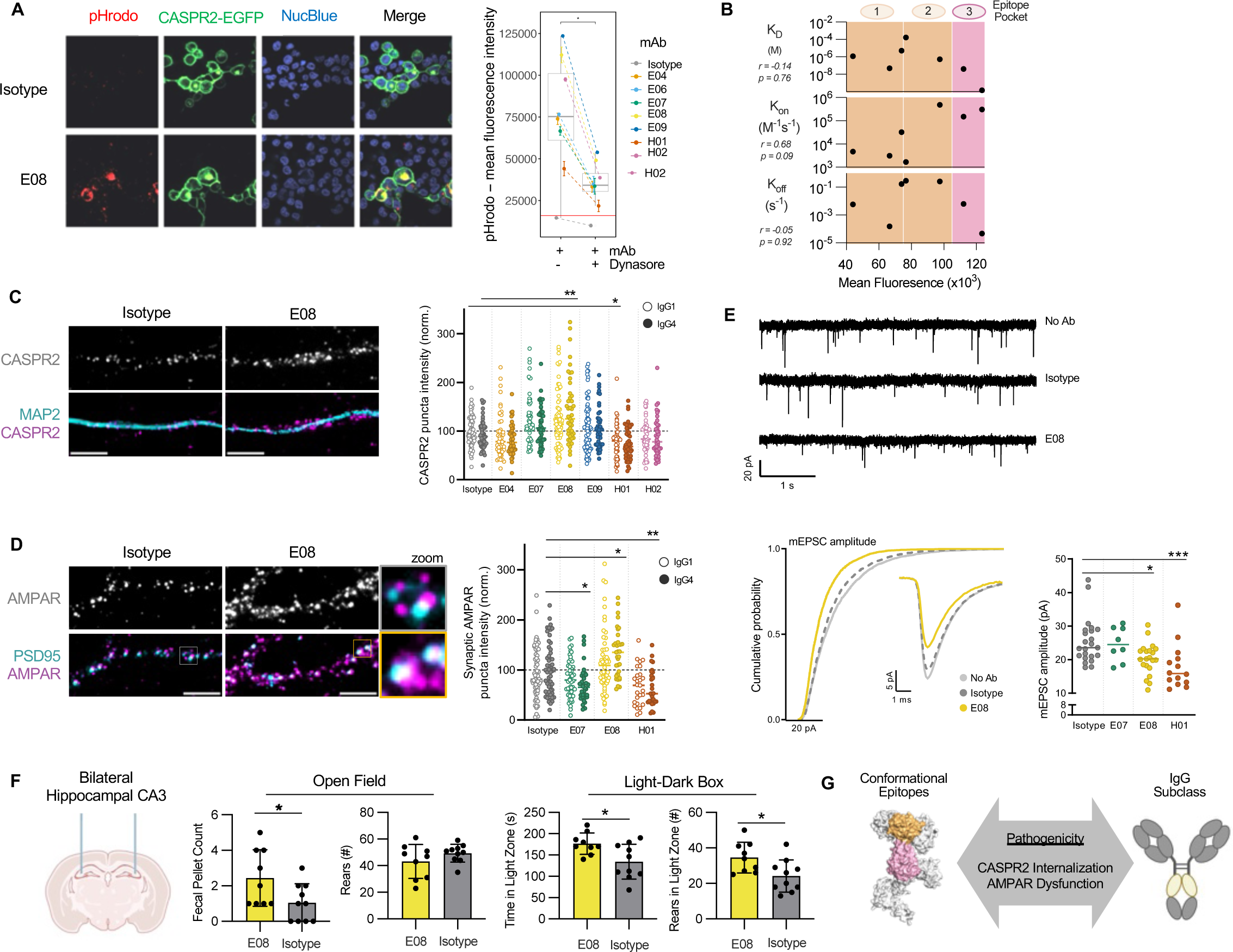
Diverse pathogenic potentials of CASPR2 mAbs. (A) Representative images used to calculate colocalisation of pHrodo-conjugated IGG4 mAbs and CASPR2-eGFP reflecting receptor internalization by CASPR2-expressing HEK293T cells (left). Internalisation quantified by mean pHrodo fluorescence intensity mAbs in the presence and absence of dynamin inhibition (Dynasore; right). (B) Pearson’s correlation of receptor internalization to mAb binding parameters; bottom x-axis label denotes internalization. Background colors indicate the 3 epitope pockets as presented in Fig.3; top x axis label annotates epitope. (C) Representative images of CASPR2 and MAP2 expression after 4 hour application of IgG1 (open circles, left) or IgG4 mAbs (closed circles); Scale bar = 5 μm. CASPR2 puncta intensity summarized on the right with *p<0.05, non-parametric Kruskal-Wallis with Dunn’s multiple comparison post-hoc test. mAbs colored as in A. (D) Representative images (left) and quantification (right) of synaptic AMPAR and PSD95 expression similar to (C). (E) Representative tracings (top), single average event and cumulative probability (middle), and amplitude quantifications (bottom) of AMPAR-mediated mEPSC recordings of pyramidal neuronal cultures. * = p<0.05, Kruskal-Wallis test with Dunn’s multiple comparison post-hoc test. (F) Cartoon depiction of intracerebral mAb injection into bilateral hippocampal CA3 regions (left). Open field (middle) and light-dark box (right) behavioral test performance was assessed at 6 and 9 hours post-injection, respectively. *=p<0.05, t-test. (G) Summary model cartoon.

To model functional effects in a more translational system, HEK293T cells were substituted for rodent neuronal cultures. CASPR2 expression, glutamatergic AMPAR expression and synaptic currents were assessed (Fig.4C-G/S3),(32) molecular alterations which may account for the seizures, amnesia and psychiatric features observed in CASPR2-Ab-E. Also, as patient CASPR2-IgGs are often IgG1 or IgG4s,(8, 13) the antibody subclass dependence of mAb effects was studied.

Two memory mAbs modulated neuronal CASPR2 expression, with dependence on IgG subclass (Fig.4C/S3): E08 upregulated CASPR2 as a IgG4 (p=0.019), whereas H01 downregulated CASPR2 as an IgG1 (p=0.002). Both mAbs not only influenced CASPR2 expression but also increased or decreased the intensity of synaptic AMPAR punctae (co-localizing with PSD95), with the same directionality as CASPR2 modulation (Fig.4D), and additionally decreased the amplitude of AMPAR-mediated miniature excitatory postsynaptic currents (mEPSCs; Fig.4E). Further highlighting their overall diversity, another mAb (E07) showed no effect on CASPR2 expression but reduced AMPAR expression as an IgG4 without an effect on mEPSCs (Fig.4C-E). As proof-of-concept towards pathogenic significance *in vivo*, E08 was stereotactically injected into the CA3 region of rat hippocampi. After only 6-9 hours, by comparison to isotype control mAbs, E08-injected rats showed increased defaecation and more time spent and rears performed in the light zone of a light-dark box (Fig.4F). These behavioral phenotypes are consistent with affective aspects of CASPR2-Ab-E patients.(8, 9)

Collectively, these functional assays show that individual CASPR2 mAbs derived from the memory compartment of patients with CASPR2-Ab-E can differentially induce rapid molecular, cellular and systems-level alterations including AMPAR dysfunction, synaptic reorganization and behavioural alterations. Effects of these disease-relevant BCRs are associated with select mAb characteristics, including epitopes and IgG subclasses (Fig.4G).

### CASPR2 naïve B cell receptor sequences contrast in health versus disease

Next, we aimed to identify BCR features facilitating escape of these disease-relevant MBC-derived mAbs from the patient NBC compartment. As NBC selection into the memory compartment likely relates to BCR signalling strength,(33), we hypothesized that naïve CASPR2-reactive BCR sequence characteristics in patients would differ from the effectively tolerised CASPR2-reactive NBCs isolated from HCs.

First, compared to anticipated VH family distributions (Fig.5A),(34, 35) all CASPR2-reactive B cell populations in health and disease favoured VH3 family use (population usage 43.1% vs 63.6% in HC (p=0.0003), 62.5% in CASPR2-Ab-E NBC and 71.4% MBCs (both p<0.0001, Chi-Squared). Despite this shared VH3 preferential usage, specific V genes and combinations of paired heavy chain V-J genes were distinct between NBCs isolated from HCs and CASPR2-Ab-E patients (Fig.5B; p<0.0001, Fisher’s exact). Only two BCRs with overlapping V-J genes were observed, and these differed in their paired light chains (Table.S2). Therefore, despite their equivalent frequencies (Fig.1C), no CASPR2-reactive NBCs across patients and HCs showed identical paired sequences. Within MBCs (whose V-J gene combinations are, by definition, identical to those in the corresponding UCAs), one V-J gene pair overlapped with the CASPR2-Ab-E NBCs (again, with a different light chain), but not with any V-J combinations from HC NBCs (Fig.5B). Moreover, affected patients preferentially used two V gene segments (VL 1-39 and 3-1), and only one light chain V-J combination overlapped between HC and CASPR2-Ab-E NBCs. In contrast, three MBC V-J combinations overlapped with the NBC population from CASPR2-Ab-E patients, but not those of HCs. More traditional correlates of autoreactivity including HCDR3 length, HCDR3 charge, and κ:λ ratio,(14, 16) showed no differences between health and disease (Fig.5C-E).

**Figure 5:**
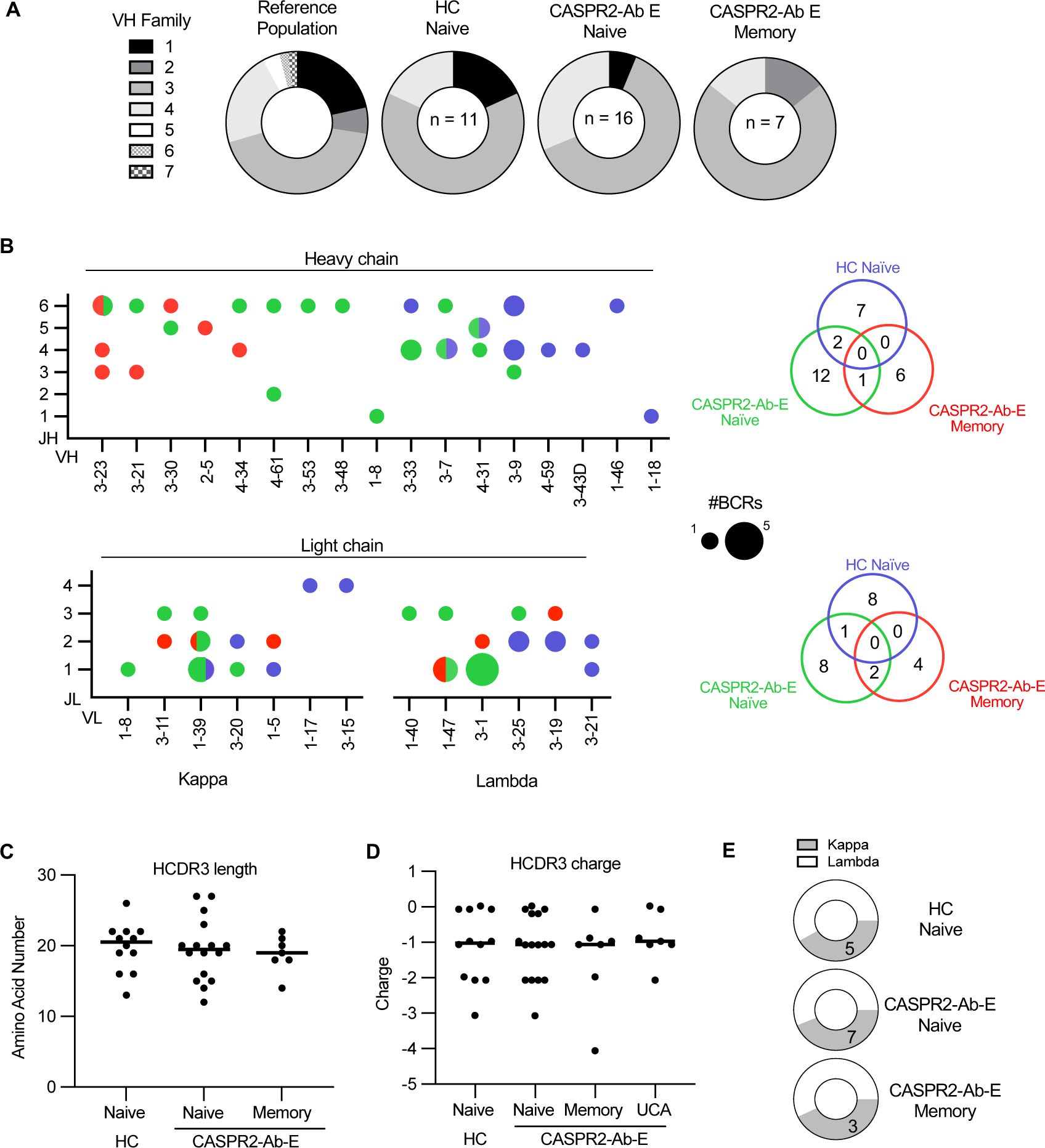
Distinct CASPR2-reactive BCRs mature in affected patients. (A) Pie charts depicting the percentage of VH family usage in each labeled population. (B) Heavy chain germline VH and JH usage (top) and light chain germline VL and JL (bottom) usage within the indicated B cell populations. Circle size corresponds to frequency. Venn diagrams (right) depict the absolute number of unique V-J pairs by population, p<0.0001, 2x2 contingency analysis comparing naïve groups. (C/D) Heavy chain CDR3 amino acid length (C) and CDR3 net charge (D) did not statistically differ (ANOVA, p>0.05). (E) Pie charts demonstrating similar ratios of kappa and lambda light chain usage by population.

Hence, fundamentally different CASPR2-reactive BCRs escape from central tolerance into the circulating NBC compartment in health versus disease, and only the naïve BCRs from CASPR2-Ab-E patients show sequence-based similarities to those from MBCs.

### Unmutated CASPR2-reactive BCRs properties contrast in health versus disease

Finally, to interrogate the fundamental basis of B cell tolerance, we hypothesized that these divergent BCR repertoires determine sequences which confer relative potentials to escape immune checkpoints. Hence, we assessed unmutated mAb binding strengths to an array of human autoantigens, in particular CASPR2, focusing on a comparison between those known to have entered the MBC compartment - the UCAs - with those *ex vivo*-derived naïve BCRs from CASPR2-Ab-E patients and HCs which likely never entered, and were not detected within, the MBC population.

First, relative avidity to CASPR2 was determined by live cell-based assay: while all NBCs bound as recombinant IgMs, none of 16 CASPR2-Ab-E-derived NBC mAbs compared to 5 of 11 healthy NBC-derived mAbs (p=0.0057, Fisher’s test) and all four CASPR2-reactive disease UCAs (p=0.0002, Fisher’s test) which bound as either IgG or Fabs (Fig.6A). This trend was also reflected in end-point dilutions: by comparison to those from CASPR2-Ab-E patients, NBC mAbs from HCs remained bound to CASPR2 at lower quantities (Fig.6B; p=0.06; Kruskal-Wallis with Dunn’s test for multiple comparisons). Most strikingly, of all three unmutated populations, the UCAs bound CASPR2 at by far the lowest observed end-point dilutions (p=0.001, vs. UCA, and p=0.04 vs. HC naive).

**Figure 6:**
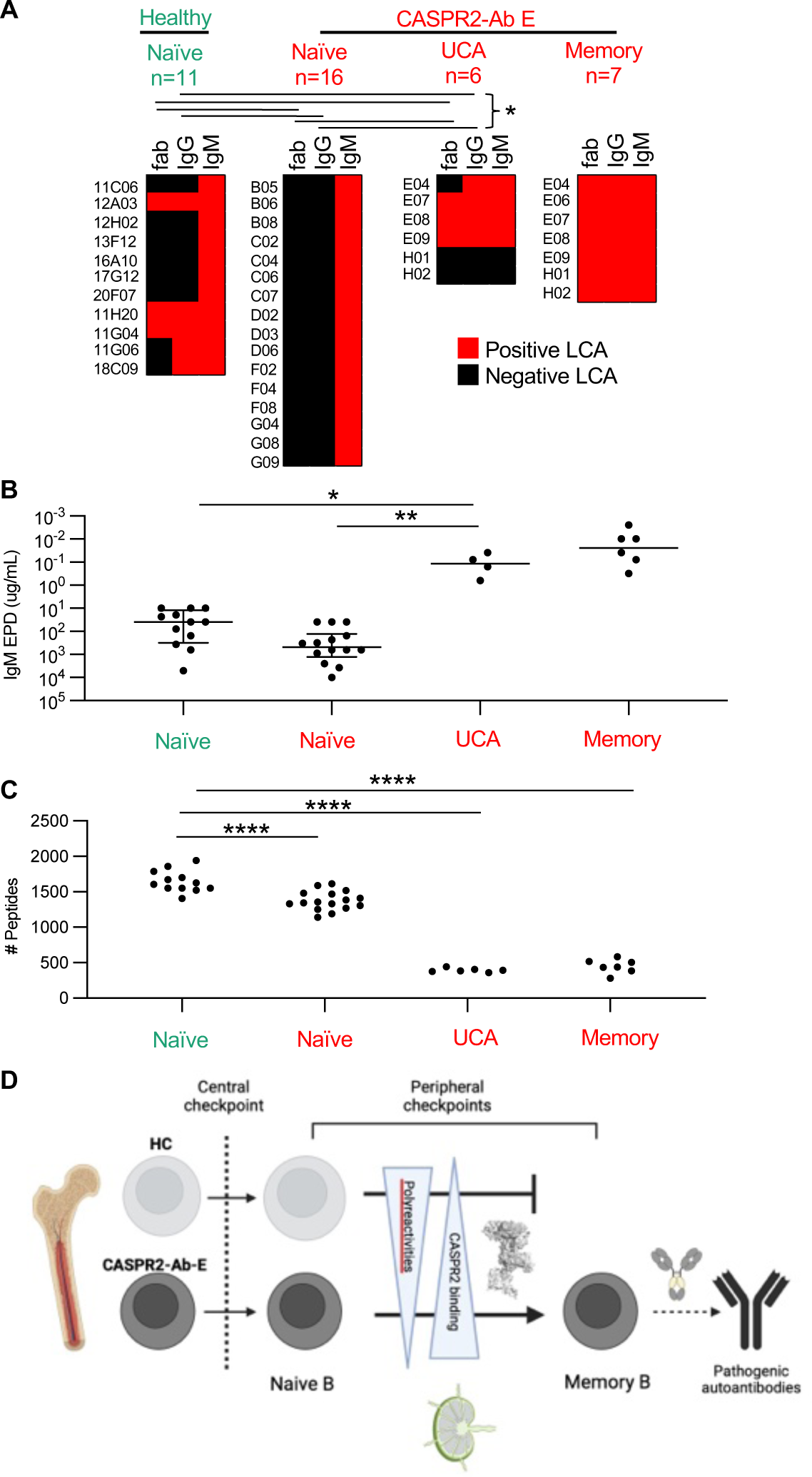
High CASPR2 avidity and otherwise low self autoreactivity facilitate clonal escape. (A) Heat maps depicting binding by CASPR2-reactive Fab segments, -IgG or -IgM, red = positive binding. * = p<0.05 for fab and IgG frequency distributions. Populations labeled in green depict healthy control, populations labeled in red depict CASPR2-Ab E throughout the figure. (B) CASPR2-IgM mAb end point dilution in a live cell based assay. All mAbs expressed as class IgM to prevent class confound * = p<0.05, ** = p<0.005 (C) Quantification of number of peptides enriched in phage immunoprecipitation, ****=p<0.0001, Wilcoxon unpaired. (D) Summary cartoon model.

Next, to mimic other autoantigens likely available to NBCs in germinal centres, we assessed broader self-reactivities using two platforms: i) a limited series of canonical autoantigens representing polyreactive (double-stranded DNA, insulin and lipopolysaccharide) or autoreactive (HEp-2 cell) antigen substrates (Fig.S4)(14) and ii) a phage-expressed array of self-antigens tiled across the entire human proteome (Fig.6C).(27) When CASPR2-Ab-E patient NBCs were compared to NBCs from HCs, rates of polyreactivity trended (3/16 (18.8%) vs 4/11 (36.4%); Fisher’s exact test p=0.39), and autoreactivity rates were statistically significantly lower (Fig.S4; 2/16 (12.5%) vs 6/11 (54.5%); Fisher’s exact test p=0.033). UCAs showed reactivities akin to the CASPR2-Ab-E patient NBCs.

More quantitatively, from the phage-display, higher rates of self-reactivities were also observed from HC NBCs compared to those from CASPR2-Ab-E patients (Fig.6C; 1657 ± 157 vs 1360 ± 141, respectively; ordinary one-way ANOVA with Tukey’s test for multiple comparisons, p<0.0001). Yet, and by contrast to CASPR2 binding strengths, the lowest self-reactivities were observed in UCAs (p=0.0001), similar to MBC characteristics.

In summary, when compared to those originating from CASPR2-Ab-E patients, the CASPR2-reactive naïve BCRs from HCs exhibited stronger binding to both CASPR2 and multiple other autoantigens. Most strikingly, UCAs showed around three-logs higher affinity for CASPR2 accompanied by very limited self-reactivity towards a comprehensive array of other human autoantigens, both similar to the MBC BCRs. These findings suggest that a balance between specifically binding CASPR2 while ignoring other autoantigens may help determine selection versus elimination of autoreactive BCRs at peripheral immune checkpoints in humans (Fig.6D).

## Discussion

This study leverages CASPR2-antibody encephalitis, a prototypical single autoantigen-directed condition, as a rare opportunity to molecularly dissect the integrity and thresholds of autoantigen-specific B cell checkpoints in both health and disease. Our observations construct a ‘multihit’ model of immunopathogenesis which begins with promiscuous early central tolerance checkpoints that release unmutated CASPR2-reactive BCRs at comparable frequencies in both health and disease states, but with different BCR sequences. UCA studies revealed that the combination of strong reactivity for native CASPR2 together with a lack of other proteome-wide self-reactivities most effectively facilitates the escape of CASPR2-reactive BCRs into the memory compartment. Although this was strictly observed in patients, memory compartment access alone was insufficient to confer pathogenicity. The rapidly-induced neuronal dysfunction appeared influenced by other BCR properties, including IgG subclass and epitope preferences. Taken together, these multimodal findings both highlight basic human immunological tolerance mechanisms and, simultaneously, aid the rational design of autoantigen-specific immunotherapies in CASPR2-Ab-E. We anticipate this comprehensive approach can be applied towards mechanistic tolerance threshold insights across all human autoantigen-specific diseases, to draw both parallels and distinctions in disease-specific characteristics.

A novel aspect of our study is the direct isolation of autoantigen-reactive NBCs. Overall, a substantial fraction of NBCs displayed CASPR2-reactivity (∼0.5%), an impressively large subset dedicated to CASPR2 given ∼20,000 non-modified proteins exist in the human proteome. This permissive central tolerance to CASPR2 was reinforced by the observation that most UCAs bound CASPR2. Further, its fundamental basis showed differences in HCs versus patients, based on immunoglobulin gene usage. Overall, the frequent tendency to early-lineage CASPR2 reactivity in association with VDJ-recombination differences suggests the immunological origins of CASPR2-Ab-E begin prior to the central B cell tolerance checkpoint. This may reflect the almost exclusive neuronally-restricted expression of CASPR2, limiting its availability for bone marrow B cell tolerization.(15)

Downstream in B cell development, the stringent deletion of bone marrow released CASPR2-reactive BCRs in HCs suggests later peripheral checkpoints effectively maintain tolerance against CASPR2 in health. However, in patients, BCRs which definitively entered the MBC compartment (studied as UCAs) appeared distinctively ‘tuned’ to balance strong native CASPR2 reactivity with minimal binding to other autoantigens. This may reflect their preferential ability to exclusively acquire CASPR2-reactive T cell help. Conversely, the broad polyreactivity combined with limited CASPR2-reactivity - as observed in most CASPR2-reactive NBCs - may confer a higher likelihood of being more readily sensed and tolerised, and hence strictly excluded from the memory compartment. This concept is consistent with rodent studies which suggest BCR affinity shapes downstream memory compartments,(36) and provides a rare mechanistic insight into the properties of human tolerance thresholds.

Within MBCs, knockout and proteome wide screens showed that CASPR2-reactivities were highly-specific for native CASPR2 conformations, and exclusive to patients. These memory BCRs showed small clonal expansions and carried mutations which conferred substantial increments in binding to human CASPR2. These observations suggest conformationally-native CASPR2 is presented in germinal centres. Peripherally, we hypothesise these reside in cervical lymph nodes,(37) which are thought to drain the CNS.(26, 38) Yet, the highest affinity mAb with the greatest internalisation capacity had acquired only five variable region mutations, not precluding a role for extrafollicular responses.(39) Nevertheless, this peripherally-dominant process of autoantibody production is consistent with the far higher levels of CASPR2-autoantibodies in serum versus cerebrospinal fluid,(2) and the relatively low rate of intrathecally-generated somatic hypermutations observed in patient cerebrospinal fluid CASPR2-reactive B cells.(40)

Yet, the entry of mutated CASPR2-reactive B cells into the MBC compartment does not appear sufficient for pathogenesis. Many MBC-derived BCRs possessed limited effects *in vitro* and, overall, showed striking diversity across multiple parameters including kinetics of CASPR2 binding and their relative capacities to induce correlates of pathogenicity: CASPR2 internalization and modulation, AMPAR clustering, and altered synaptic kinetics. Also, their pathogenicity in individual assays varied based on subclass and epitope. Hence, we propose that only some mAbs confer direct pathogenic potential. This conclusion may explain why overall levels of serum CASPR2-IgG do not correlate with disease status, and help reconcile functional differences between previous experiments which studied polyclonal serum IgGs.(30, 31) By extension, we hypothesise that a detailed deconvolution of mAb properties within polyclonal samples may more accurately correlate with clinical and functional features. Further, these observed functional molecular alterations offer future targets to potentially modify symptoms of CASPR2-Ab-E.

An important outstanding question raised by our model is whether the observed loss of both central and peripheral tolerance is continually present in affected patients, or instead represents a ‘one-time’ dysregulation. Probing checkpoint function longitudinally may provide further insights and guide therapeutic strategies that not only limit the potential for adverse effects, but also inform the natural history of the disorder. These could be complemented with measures of germinal centre activity, such as CXCL13 levels and antigen-specific IgMs,(26, 38) to redefine composite prognostic biomarkers. Such approaches may improve relapse prediction and guide preventative therapeutic strategies.

Additionally, our observations help rationally direct future therapeutic considerations. For example, restoration of peripheral tolerance with regulatory T cells may represent a logical approach to effectively treat CASPR2-Ab-E, as these cells can limit autoreactive B cell accumulation.(41) Central tolerance defects may also be combated using approaches that modulate this checkpoint.(15) As elimination of both CASPR2-reactive memory and naïve B cells may be required for long-lasting effective treatment, broad B cell lineage depletion (for example with CD19 rather than CD20 directed therapeutic antibodies) may hold most promise.(42) Also, our neurobiological findings inform more downstream therapeutics by highlighting AMPAR upregulation as a goal for symptomatic benefit.

Our data also speak to diagnostic paradigms. The high frequencies of naïve CASPR2-reactive BCRs may explain the observed false-positive rates of serum CASPR2 antibodies.(24, 43) Further, to reduce false negative results, differences in cross-species reactivities should encourage the use of human based substrates in diagnostic laboratories, rather than rodent brain sections and cultured neurons.(44)

Limitations of our work include the low numbers of patients and HCs studied. Yet, our focus was on the detailed characterisation of >12,000 B cells and 37 mAbs from untreated patients, who are difficult to recruit in a rare disease. Also, we did not examine B cells from cerebrospinal fluid or cervical lymph nodes, sites closest to the neuronal dysfunction.(26, 38, 40, 45) Nevertheless, our observations suggest substantial affinity maturation and loss of tolerance begin in the periphery, consistent with independent observations from cerebrospinal fluid.(40) Also, the absence of clonal overlaps between NBC and MBC compartments necessitated use of UCAs to study the evolution of individual BCRs. While UCAs are widely used to infer germline characteristics, they may fundamentally fail to capture the junctional diversity observed in native NBCs.(20) Finally, our experiments did not examine factors responsible for a faulty peripheral checkpoint in patients, such as polymorphisms in key checkpoint molecules and the presence of CASPR2-reactive T cells.(46) Future experiments should study these in addition to the relative importance of other end-effector mechanisms including complement fixation, FcR binding and Fab-Fab arm exchange.

In summary, our data reveal key aspects of the therapeutically-tractable immunobiology and neurobiology underlying CASPR2-Ab-E and present a novel roadmap to systematically dissect how sequential, comparative studies of autoantigen-specific B cell inform our understanding of how and where immunological tolerance is lost in human autoimmune diseases.

## Methods

### Sex as a biologic variable

Our study examined male and female humans. Investigators were blinded to the sex of some healthy controls precluding further generalizations.

### Ethics

All human investigations were reviewed and approved by the University of Oxford, ethical approvals REC16/YH/0013 and REC16/ES/0048. All experiments involving animals were reviewed and approved under project license P996B4A4E and personal license I11739608 at the University of Oxford, or Orgão de Bem-Estar e Ética Animal (ORBEA) and Direcção Geral da Alimentação e Veterinária (DGAV) at the University of Coimbra.

### Participants and samples

CASPR2-antibody encephalitis patients (n=6) and healthy participants (n=5) provided written informed consent. Available clinical information is summarized in Table S1; researchers were completely anonymized to further details of three of the healthy controls. Donated peripheral blood mononuclear cells (PBMCs) were cryopreserved in liquid nitrogen until use.

### Fluorescence activated cell sorting with bulk and single cell lymphocyte cultures

PBMCs were thawed, labelled with antibodies against CD3, CD14, CD19, CD27 and IgD, and subsequently fluorescence-activated cell sorted for naïve B cells (CD19^+^IgD^+^CD27^-^) and class-switched memory B cells (CD19^+^IgD^-^CD27^+^) from CD3^-^CD14^-^DAPI^-^ lymphocytes. A complete list of all primary and secondary antibodies used in this study is provided in Table S5.

Conditions for 13-day bulk(22, 38) and 22-day single cell cultures(26) are reported. In brief, for bulk cultures 10,000 of each B cell subset were sorted and cultured in complete B cell media (RPMI with 5% IgG depleted foetal calf serum) with R848 (2.5 µg/ml, Enzo Life Sciences), soluble CD40L (50ng/ml, R&D systems), interleukin-2 (50 ng/ml Peprotech), interleukin-1β (1ng/ml), interleukin-21 (50 ng/ml PeproTech), interleukin-6 (10 ng/ml R&D Systems) and tumour necrosis factor-a (1 ng/ml PeproTech). For single cell cultures, single sorted B cells were cultured for 22 days with MS40L^low^ cells (kindly gift from Garnett Kelsoe), (1, 2) supplemented with interleukins-2, -21 and -4.

### Culture supernatant screening and live cell-based assays

Well-described live cell-based assays(5, 6) were used to test for the presence of CASPR2-reactive IgM/IgGs in 50 µl of culture supernatant, or quantify mAb antigen-specificity and end-point dilutions (starting at 10 µg/ml). In brief, live HEK293T cells transiently transfected with either a human or mouse CASPR2-EGFP intracellular C-terminus fusion construct were exposed to supernatant or mAbs for 45 minutes, and subsequently washed and fixed. Bound antibodies were detected with secondary antibodies against either human-IgG or -IgM.

### Generation of recombinant CASPR2-reactive antibodies

RNAsin plus (12.8ul/ml, Promega) in TE buffer pH8 (Bioultra) was added to CASPR2-antibody positive wells and flash frozen. Single cell RNA was made into cDNA from which heavy and light chain immunoglobulin variable region genes were amplified as described (Table S4),(47) and validated with Sanger sequencing. Sequence annotation and analyses (including germline VDJ usage, mutation count, HCDR3 length and charge) were performed using IMGT High-Vquest (which reports established polymorphisms) and spectral clustering for clone partitioning. Validated variable domain genes were then cloned into expression vectors containing mu, gamma 1, or gamma 4 constant-region Ig domains, or hexahis-tagged FAB fragments.

For production, HEK293F cells cultured in Freestyle expression medium (Thermofisher) were PEI-transfected with vectors encoding cognate-paired heavy and light chain sequences, at a 1:1 vector ratio. For IgG1/IgG4 mAbs, supernatant was harvested after 5 days and IgGs purified on a single AKTA pure chromatography system. For IgM mAbs, a J-chain expression plasmid was co-transfected and IgMs purified using centrifugal size exclusion. Hexahis-tagged FAB fragments were purified using nickel affinity chromatography.

### UCA reversion

Somatically hypermutated variable region sequences were aligned with IMGT-VQUEST to identify the ancestral germline gene fragments (https://www.imgt.org). Replacement mutations were reverted to the respective nucleotides from the best matched germline gene. Non-templated nucleotides at V-D, D-J and VL-JL junctions were unmodified. Finally, using the IMGT junctional tool, CDR3 regions of the heavy chains were also back-mutated to the best aligned D-gene. Reverted sequences were ordered as gene fragments (IDT) with flanking cloning sites and inserted into IgG, IgM and FAB expression vectors.

### Surface plasmon resonance

A BIAcore 8k (Cytiva) measured affinity (*K_D_*), and kinetics (association and dissociation constants: *k_on_* and *k_off_*) of immobilised mAb binding to the C-terminally biotinylated purified extracellular domain of human CASPR2. Approximately 200 response units (RU) of IgG was captured onto a Protein A Series S Sensor Chip (Cytiva). Single cycle kinetic analysis of binding was undertaken, using two-fold serial dilutions of CASPR2 in HBS-P+ buffer (Cytiva) and six dilutions per cycle. The CASPR2 starting concentrations varied from 20 nM to 1000 nM. All measurements were made with injection times of 120s (30 µl/min) and dissociation times of 600s (30 µl/ml) at 37°C. Regeneration of the sensor chip was performed with 10 mM Glycine-HCl, pH 1.7 for 30s (30ul/min) between experiments. For analysis, the sensograms were double reference subtracted and fitted with a 1:1 binding model using the Biacore Insight Evaluation Software version 2.0.15.12933 (Cytiva). For all 1:1 binding fits the t_c_ value was set at 1e+10.

### Antibody binding to self and non-self substrates

Immunoblot analysis was performed per manufacturer’s instruction of electrophoresis and western blotting system (Life technologies). Briefly, purified extracellular domain of CASPR2 or HEK cell lysate (generated by adding RIPA buffer containing protease inhibitor cocktail over ice, Thermo Fisher) were protein sources. CASPR2 protein was denatured by LDS buffer containing beta-mercaptoethanol. Samples were separated by 10% Bis-Tris precast gels, and transferred to polyvinylidene fluoride (PVDF) membrane. Nonspecific binding was blocked by 5% skim milk (Sigma) in TBS/0.05% Tween-20 (Life technologies) at room temperature. The blot was probed with antibodies of interest at 4℃ overnight. Commercial polyclonal rabbit anti-human CASPR2 antibody (Novus, A45565) was used as isotype control for detecting denatured CASPR2. Horseradish peroxidase (HRP)-conjugated secondary antibodies (Dako) were used for subsequent incubation. ECL solution (Thermo Fisher) was used for chemiluminescence. Signals were detected by ChemiDoc MP imaging system (BioRad). The exposure time was determined automatically when saturated signal was detected.

Methods to determine mAb binding to neuronal substrates (starting dilution 20 µg/ml) expressing CASPR2 have been described previously: in brief, 10 µM thick rat brain sections lightly fixed in 4% paraformaldehyde and stained with diaminobenzidine,(6) live rat dorsal root ganglion neurons,(10) live primary cultured rat hippocampal neurons,(5, 40) 4% sucrose/paraformaldehyde-fixed rat cortical neurons (32) and live human sensory-neuron IPSCs myelinated by rat Schwann cells.(48)

Polyreactivity was assessed against 20 µg/ml double-stranded DNA, 10 µg/ml lipopolysaccharide, or 15 µg/ml recombinant human insulin (all SIGMA-Aldrich), as described.(14) The highly-polyreactive antibody ED38 served as positive control. For the Hep-2 ELISA, mAbs were applied to human epithelial type 2 (HEp-2; HELA cells) cell lysates (INOVA ELISA kit) and developed with HRP (Bio-Rad). Plates were read out with the EPOCH (BIO-TEK) and positivity designated >OD of 0.7 (405 nm).

Programmable phage immunoprecipitation and sequencing (PhipSeq) investigated the conformational nature of CASPR2 epitopes as well as broader self-reactive potential. In brief, IgG1 mAbs were incubated with 10^10^ plaque-forming units/ml of a phage-display library composed of overlapping 49-mer peptides arrayed across the human proteome or CASPR2-protein. Antibody-bound phage particles were isolated by protein G immunoprecipitation and underwent MiSeq next generation sequencing to identify putative human antigens.(27)

### CASPR2-CASPR4 domain swaps and epitope binning

Gene sequences corresponding to CASPR4 protein domains discoidin, laminin G-like domains 1-4, and fibrinogen C-terminal, were identified using https://www.uniprot.org. Complementary overhangs matching flanking 5’ and 3’ regions of CASPR2 were added. CASPR4 domain ‘knock-in’ constructs were generated using these full gene constructs (IDT) via overlap extension PCR. The constructs were used in live cell assays as above.

For competitive binding experiments, CASPR2-expressing HEK293T cells were first saturated with an excess of a single unlabelled mAb (100 µg/ml). After 60 minutes, mAbs (10 µg/ml; expressed as IgG1s to avoid potential Fab-arm exchange) pre-conjugated with AF594 (Thermofisher #A20185) were added for 30 minutes incubation. Wells were washed and AF594 fluorescence detected on a BMG Omega Fluostar fluorescence plate reader (excitation 560nm, detection 610nm). Percentage inhibition was defined as the percentage reduction from maximal binding.

### Cross-species protein structures

CASPR2 mouse and human protein sequences were obtained from uniport.org. The predicted crystal structure of CASPR2 (structure ID 5Y4M, AlphaFold.ebi.ac.uk) was modeled using PyMol.

### pHrodo internalisation and image quantification

pHrodo conjugated mAbs (20 µg/ml) were incubated overnight with CASPR2-transfected HEK293T cells and mean fluorescence quantified at regular intervals using BMG OMEGA FLUOstar. For some wells, pre-treatment with 50 μM dynasore was performed 1 hour prior to mAb incubation. pHrodo mean fluorescence quantification was pre-processed in FIJI and analysed in R, using packages rstatix (https://github.com/kassambara/rstatix) and RKcolocal (https://github.com/lakerwsl/RKColocal). In FIJI, 12 regions of interest were selected and the mean fluorescence intensities for each region was exported for analysis in R.

### Primary cultures of cortical neurons

Primary cultures of rat cortical neurons were prepared from the cortices of E17/18 Wistar rat embryos, as previously described (Fernandes et al., 2019). Briefly, after dissection, tissue was treated for 10 min at 37°C with trypsin (0.06%, Gibco Invitrogen), in Ca^2+^- and Mg^2+^-free Hank’s balanced salt solution (HBSS: 5.36 mM KCl, 0.44 mM KH_2_PO_4_, 137 mM NaCl, 4.16 mM NaHCO_3_, 0.34 mM Na_2_HPO_4_.2H_2_O, 5 mM glucose, 1 mM sodium pyruvate, 10 mM HEPES and 0.001% phenol red). Cells were then washed 6 times in HBSS and mechanically dissociated. Cells were plated in neuronal plating medium (MEM supplemented with 10% horse serum, 0.6% glucose and 1 mM pyruvic acid) onto poly-D-lysine-coated (0.1 mg/mL) coverslips in 60 mm culture dishes, at the desired density. For imaging purposes, cells were plated at a final density of 2.5 x 10^5^ cells/dish; for electrophysiology experiments, cells were plated at a density of 11 x 10^5^ cells/dish. After 2-4 h, coverslips were flipped over an astroglial feeder layer in Neurobasal medium [NBM, supplemented with SM1 neuronal supplement (StemCell Technologies, Grenoble, France), 0.5 mM glutamine and 0.12 mg/ml gentamycin]. Wax dots on the neuronal side of the coverslips allowed the physical separation of neurons from the glia, despite neurons growing face down over the feeder layer. To further prevent glia overgrowth, neuron cultures were treated with 10 μM of 5-Fluoro-2’-deoxyuridine (Sigma) after 3 DIV. All cultures were maintained at 37°C in a humidified incubator of 5% CO_2_ / 95% air, until DIV14.

### Immunocytochemistry, cell imaging and quantitative fluorescence analysis

Primary cortical neurons at DIV14 were incubated for 2 h at 37°C with 20 µg/mL of each monoclonal antibody. Cortical cells were then fixed for 15 min in 4% sucrose / 4% paraformaldehyde in phosphate buffered saline (PBS – 137 mM NaCl, 2.7 mM KCl, 10 mM Na_2_HPO_4_, 1.8 mM KH_2_PO_4_, pH 7.4), permeabilized with 0.25% Triton X-100 in PBS for 5 min, and then incubated in 10% (w/v) BSA in PBS for 30 min, at 37°C, to block nonspecific staining. Cells were then incubated with primary antibodies against Caspr2, the neuronal dendritic marker MAP2 or the glutamatergic synapse marker PSD95, diluted in 3% BSA in PBS (2 h, 37°C or overnight, 4°C). Following several PBS washes, cells were incubated with the appropriate fluorophore-conjugated secondary antibodies (1 h, 37°C), and coverslips were finally washed and mounted using fluorescent mounting medium from DAKO (Glostrup, Germany). To label cell surface AMPAR subunits, live neurons were incubated for 10 min at room temperature with a pan-antibody against extracellular epitopes in the N-terminus of the GluA1 and GluA2 subunits, diluted in conditioned neuronal culture medium. Coverslips were then fixed and probed as described above.

Sets of cells that were cultured and stained simultaneously were imaged using identical acquisition settings on a Zeiss Axiovert 200M microscope with a 63 X 1.4 numerical aperture oil objective. Blind-to-condition quantification was performed in the image analysis software FIJI using an in-house developed macro to automatize quantification steps. The region of interest was randomly selected avoiding primary dendrites, and dendritic length was measured using MAP2 staining. Surface AMPAR and Caspr2 digital images were thresholded such that recognizable clusters were included in the analysis, and measured for cluster intensity, number, and area for the selected region. Synaptic clusters of AMPAR were selected by their overlap with thresholded and dilated PSD95 signal. Measurements were performed in a minimum of 3 independent experiments, and at least 10 cells per condition were analysed for each preparation.

### Electrophysiology

Whole-cell patch-clamp recordings in voltage-clamp configuration were measured from 14 DIV cortical neurons plated on coverslips following 2 h incubation with each mAb. The recording chamber was mounted in a fixed-stage inverted microscope (Zeiss Observer.A1) and perfused at a constant rate (2–3 mL/min) with extracellular solution (ECS - 140 mM NaCl, 2.4 mM KCl, 10 mM HEPES, 10mM glucose, 4 mM CaCl_2_, 4 mM MgCl_2_, pH 7.3, 300-310 mOsm), at room temperature (∼23°C). AMPAR-mediated miniature excitatory postsynaptic currents (mEPSCs) were pharmacologically isolated by adding 1 µM tetrodotoxin (TTX), 100 µM picrotoxin (PTX) and 50 µM (2R)-amino-5-phosphonovaleric acid (D-APV) to the ECS. Neurons were patched using a borosilicate glass recording pipette (tip resistance 3–5 MΩ) filled with a Cs-based internal solution (107 mM CsMeSO_3_, 10 mM CsCl, 3.7 mM NaCl, 5 mM TEA-Cl, 0.2 mM EGTA, 20 mM HEPES, 4 mM ATP magnesium salt, 0.3 mM GTP sodium salt; pH 7.3, 295–300 mOsm), and recordings were initiated 2–3 min after break-in. The EPC 10 USB patch-clamp amplifier (HEKA Elektronik) was used for voltage-clamp recordings. Cells were held at -70 mV and mEPSCs were recorded over a period of 5 min in a gap-free acquisition mode. Data was digitized at 25 kHz and acquired using the PatchMaster software (HEKA Elektronik), with a signal filter of 2.9 kHz. The Clampfit software (Axon Instruments) was used to analyse the acquired mEPSCs, using a template search method to detect events, as previously described (Caldeira, Inacio et al., 2022). The template was generated by averaging approximately 30 events and the template match threshold was set to 4. Recordings were excluded from analysis if the series resistance (R_S_) was > 25 MΩ, the holding current was > 250 pA, or if the R_S_ or holding current changed more than 20%. One hundred and fifty consecutive events that met these criteria were analysed from each cell.

### Intracerebral mAb injection

5-week old female C57BL/6J mice (Charles River) were housed in cages of five in a room maintained at a controlled temperature (21°C) and humidity (5-10%) with illumination at 12h cycles; food and water were available *ad libitum*. All injections were performed during the light phase, and animals were habituated to the experimental room for one hour prior to beginning injections and 15 minutes prior to behavioural testing. Procedures were conducted in compliance with the Animal Scientific Procedures Act (ASPA) 1986, revised in 2012, and the European Directive 63/2010 on the protection of animals, as well as in accordance with the Institutional Animal Care and Use Committee (University of Oxford).

On the day of surgery, mice were anaesthetised with isoflurane and placed in a stereotactic apparatus. A mid-sagittal incision was made to expose the cranium and two burr holes were drilled over the hippocampi to the following coordinates from bregma: anteroposterior -1.5 mm; lateral ±1.8 mm. A glass microcapillary containing the solution to be injected (E08 IgG4 mAb or an isotype control mAb) was lowered 1.5 mm ventral to bregma and 1 μl of mAb at a concentration of 1 mg/ml was injected per hemisphere. The incision was then cleaned and closed with sutures. 10 mice were injected with the Caspr2 mAb and 10 with an isotype control mAb. Mice were allowed to recover post-surgery prior to behavioural testing. Open Field test was performed 6 hours post-surgery, and Light-Dark Box test 9 hours post-surgery. Behavior was manually scored after video review.

### Open access

For the purpose of Open Access, the author has applied a CC BY public copyright licence to any Author Accepted Manuscript (AAM) version arising from this submission.

## Data Availability

All data are available in the main text, supplementary materials, or available to researchers once data/material transfer agreements are in place.

## Author Contributions

BS: acquiring and analyzing data, writing the manuscript, study design; DF: acquiring and analyzing data, study design; AK: acquiring and analyzing data, study design; SP: acquiring and analyzing data; RH: acquiring and analyzing data; SR: acquiring and analyzing data; AH: acquiring and analyzing data, writing the manuscript; MM: acquiring and analyzing data; MF: acquiring and analyzing data; RD: acquiring and analyzing data; DA: acquiring and analyzing data; HB: acquiring and analyzing data; MI: acquiring and analyzing data; RW: acquiring and analyzing data; AV: acquiring and analyzing data; ST: acquiring and analyzing data; AF: acquiring and analyzing data; RS: acquiring and analyzing data; HF: acquiring and analyzing data; VM: acquiring and analyzing data; AD: acquiring and analyzing data; MT: acquiring and analyzing data; AH: data analysis and interpretation; MK: acquiring and analyzing data; MZ: acquiring and analyzing data; JB: data analysis and interpretation; RBR: data analysis and interpretation; JP: data analysis and interpretation; JP: RD: acquiring and analyzing the data; BA: data analysis and interpretation; LD: data analysis and interpretation; SR: data analysis and interpretation; RO: data analysis and interpretation; DA: data analysis and interpretation; DB: data analysis and interpretation; PW: data analysis and interpretation, study design; SD: data analysis and interpretation; MW: data analysis and interpretation; KO: data analysis and interpretation; JS: acquiring and analyzing the data, writing the manuscript; ALC: data analysis and interpretation; SI: data analysis and interpretation, study design, writing the manuscript. Co-first authors were assigned authorship order based on time of contribution.

## Acknowledgements

This research was funded in whole or in part by (to SRI) a senior clinical fellowship from the Medical Research Council [MR/V007173/1], Wellcome Trust Fellowship [104079/Z/14/Z], the National Institute for Health Research (NIHR) Oxford Biomedical Research Centre (BRC) (The views expressed are those of the author(s) and not necessarily those of the NHS, the NIHR or the Department of Health), an Association of British Neurologist Clinical Research Training Fellowship via the Patrick Berthoud Charitable Trust [2018-PBCT-1; to BS], R01MH122471 and Westridge Foundation grants (to MRW), a DFG Research fellowship (FI 2471/1-1; to MLF); COMPETE 2020 - Operational Programme for Competitiveness and Internationalisation and Portuguese national funds via FCT – Fundação para a Ciência e a Tecnologia, under project[s] POCI-01-0145-FEDER-029452 and UIDB/04539/2020, UIDP/04539/2020 and LA/P/0058/2020 (to ALC).

## Notes

### Competing Interest Statement

SRI has received honoraria/research support from UCB, Immunovant, MedImmun, Roche, Janssen, Cerebral therapeutics, ADC therapeutics, Brain, CSL Behring, and ONO Pharma, and receives licensed royalties on patent application WO/2010/046716 entitled Neurological Autoimmune Disorders. And has filed two other patents entitled Diagnostic method and therapy (WO2019211633 and US-2021-0071249-A1; PCT application WO202189788A1) and Biomarkers (PCT/GB2022/050614 and WO202189788A1). MRW receives unrelated research grant funding from Roche/Genentech and Novartis, and received speaking honoraria from Genentech, Takeda, WebMD and Novartis. KCO is an equity shareholder of Cabaletta Bio. MLF has received speaker's honoraria by Alexion, received a SPIN award from Grifols (outside the submitted work) and is a member of the Alexion-Akademie since 2022. DLB has acted as a consultant for 5 am ventures, AditumBio, Astra Zeneca Biogen Biointervene, Combigene, LatigoBio, GSK, Ionis, Lexicon therapeutics, Neuvati, Novo Ventures, Olipass, Orion, Replay, SC Health Managers, Third Rock ventures, Vida Ventures, Vertex on behalf of Oxford University Innovation. AJD is named inventor on patent pending: Immune cell therapy for nerve damage (WO2020009437A1, US-2021-0121501-A1).

### Summary of Updates

This version of the manuscript has been revised to update the following: abstract, results, and figures are updated to represent the original manuscript file

