## Supplementary material for "Permissive central tolerance plus defective peripheral checkpoints licence pathogenic memory B cells in CASPR2-antibody encephalitis": Data Supplement

Bo Sun et al.

Contents:

Figures S1-4

Tables T1-5

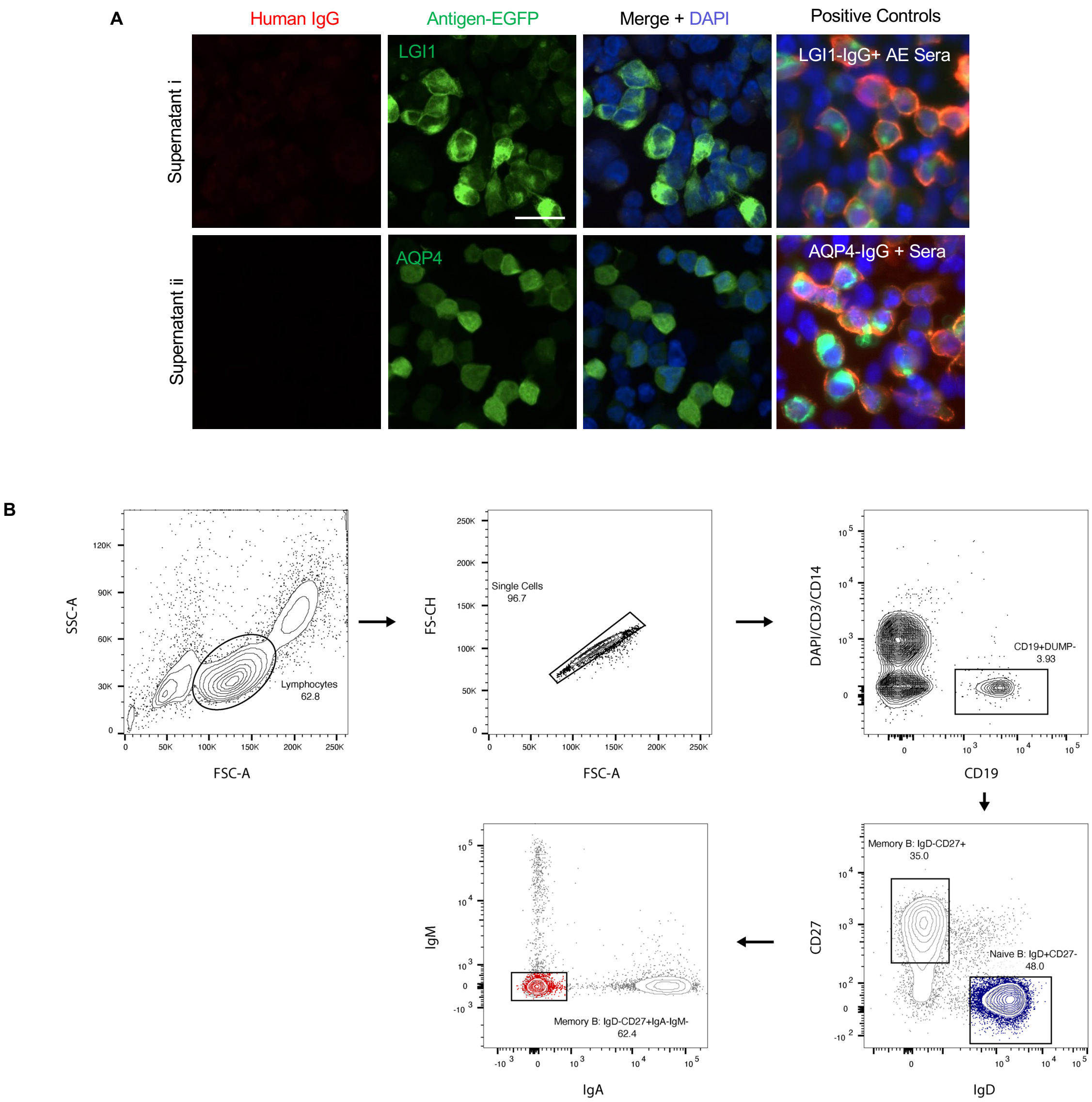


**Figure S1: B cell sorting and supernatant screening**

(A) Representative images demonstrating that bulk culture supernatants with CASPR2-reactivities showed no binding to other neuronal autoantigens, including LGI1 and AQP4. Positive control serum from an LGI1 antibody-positive patient and an AQP4-antibody positive patient are depicted (right). Scale bar = 1 micron. (B) Flow cytometry gating strategy for single cell cultures.

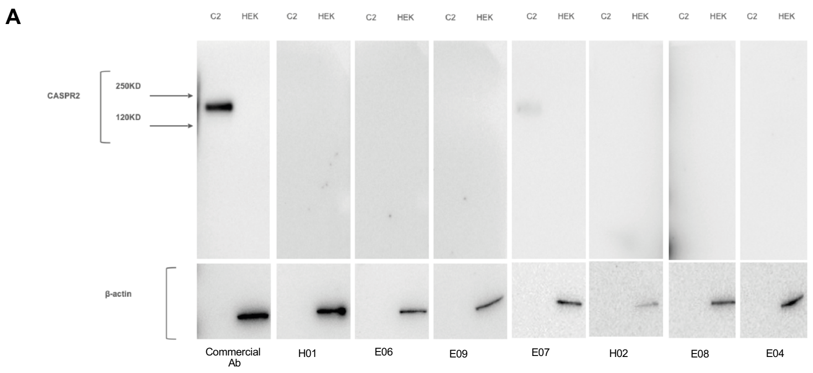


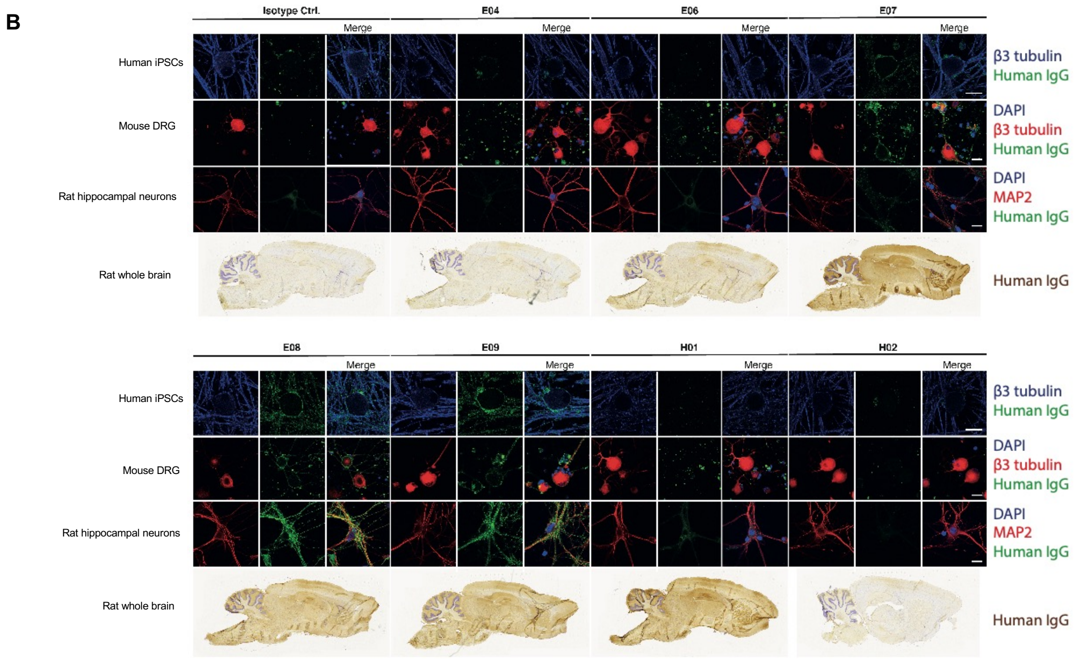


C

| CASPR2-Ab E | B Cell Subset | Human iPSC Sensory Neuron | Rodent DRG | Rodent Hippocampal | Rodent Brain |
|---|---|---|---|---|---|
| B05 | Naïve | | - | - | - |
| B06 | Naïve | | - | - | - |
| B08 | Naïve | | - | - | - |
| C02 | Naïve | | - | - | - |
| C04 | Naïve | | - | - | - |
| C06 | Naïve | | - | - | - |
| C07 | Naïve | | - | - | - |
| D02 | Naïve | | - | - | - |
| D03 | Naïve | | - | - | - |
| D06 | Naïve | | - | - | - |
| F02 | Naïve | | - | - | - |
| F04 | Naïve | | - | - | - |
| F08 | Naïve | | - | - | - |
| G04 | Naïve | | - | - | - |
| G08 | Naïve | | - | - | - |
| G09 | Naïve | | - | - | - |
| E06 | Memory | + | - | - | - |
| H02 | Memory | + | - | - | - |
| E04 | Memory | + | - | - | - |
| H01 | Memory | + | + | - | - |
| E07 | Memory | + | + | + | + |
| E08 | Memory | + | + | + | + |
| E09 | Memory | + | + | + | + |

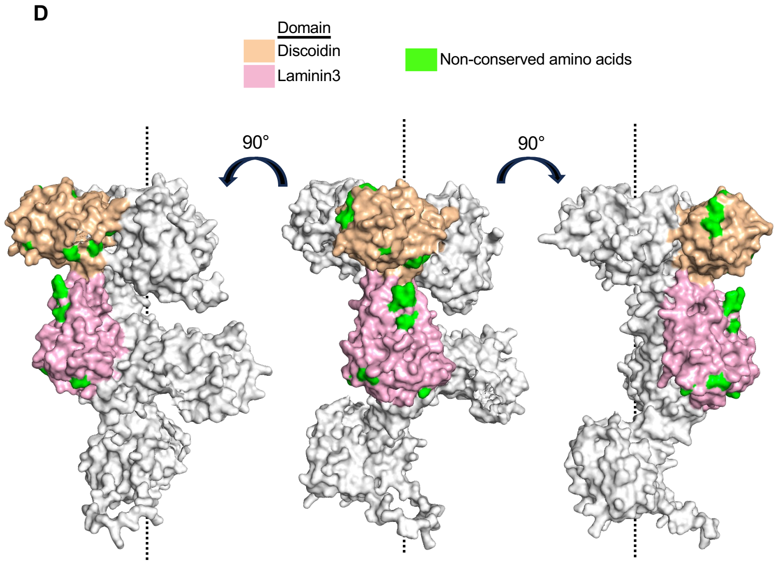


**Figure S2: mAb binding to conformational epitopes**

(A) Representative western blot of mAb binding to purified CASPR2 protein (C2) or HEK cell lysate (HEK). Beta-actin as protein loading control. (B) Representative images of CASPR2 mAbs binding to live iPSC-derived human sensory neurons (first row; identified with Beta-tubulin III in blue), live mouse primary dorsal root ganglia (DRG; second row; Beta-tubulin III in red), live primary rat hippocampal neurons (third row; identified with microtubule associated protein 2, MAP2, red) and paraformaldehyde-fixed sagittal sections of rat whole brains. Binding patterns are summmarised in (C). (D) The predicted CASPR2 protein structure is rotated along the depicted axis to better visualize the entirety of the discoidin (tan) and laminin 3 (pink) domains. Surface exposed non-conserved amino acids are depicted in green.

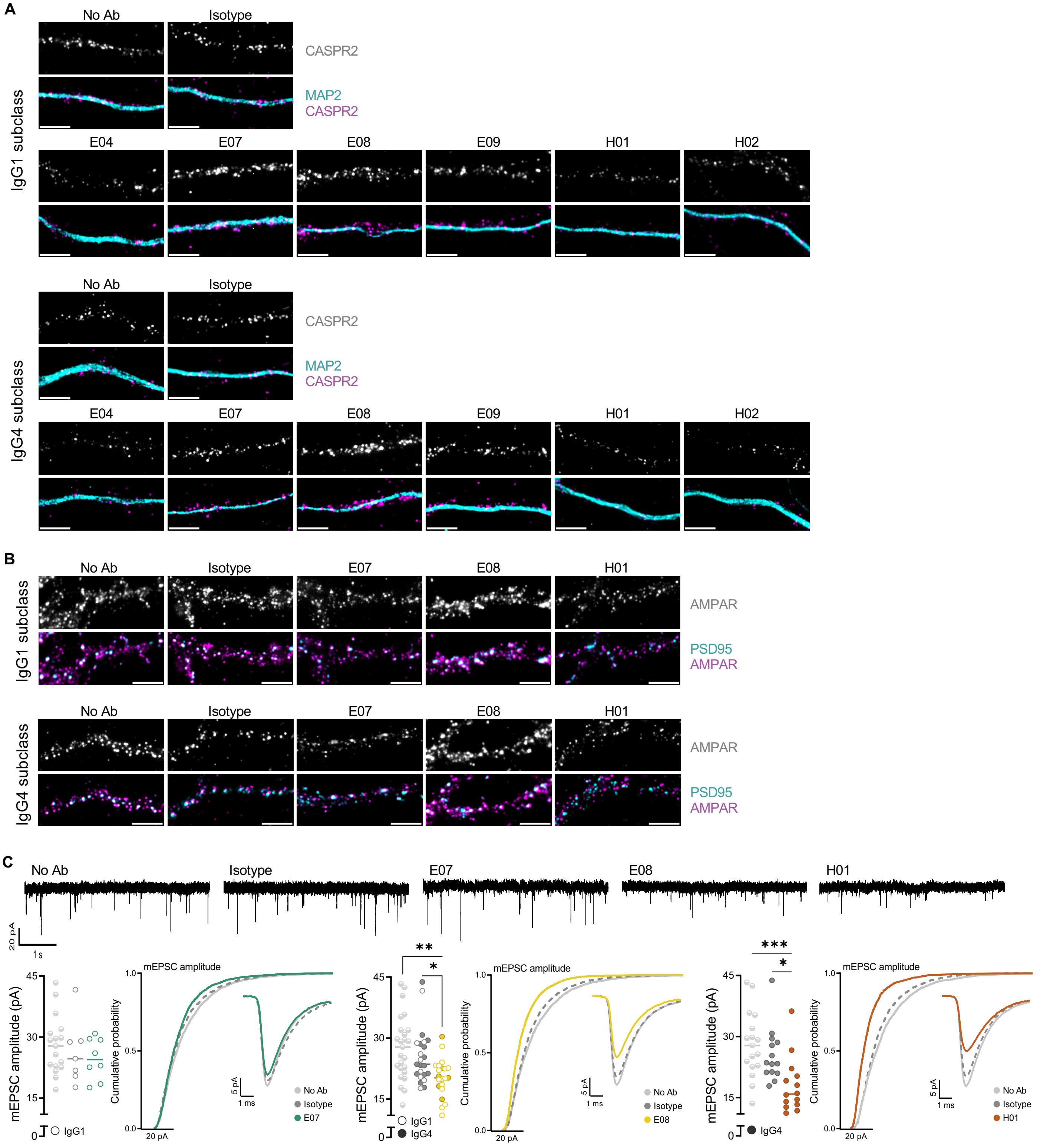


**Figure S3. Diverse effects of CASPR2 mAbs on CASPR2 and AMPAR expression, and synaptic currents.**

(A) Representative images of CASPR2 and MAP2 expression after 2 hour application of the various IgG1 (top) or IgG4 mAbs (bottom) quantified in Fig.4C; Scale bar = 5 µm. (B) Representative images of synaptic AMPAR and PSD95 expression following incubation of the E07, E08 and H01 mAbs in either IgG1 (top) or IgG4 (bottom) subclass, as quantified in Fig.4D. (C) Representative tracings (top) of AMPAR-mediated mEPSC recordings of pyramidal neurons following incubation of the E07, E08 and H01 mAbs in different subclasses, pooled together in Fig.4E. (Bottom) Average amplitude quantifications, cumulative probability and single average event of cells incubated with E07 (left), E08 (middle) or H01 (right) mAbs. *=$p<0.05$, **=$p<0.01$ and ***=$p<0.001$, Kruskal-Wallis test with Dunn's multiple comparison post-hoc test.

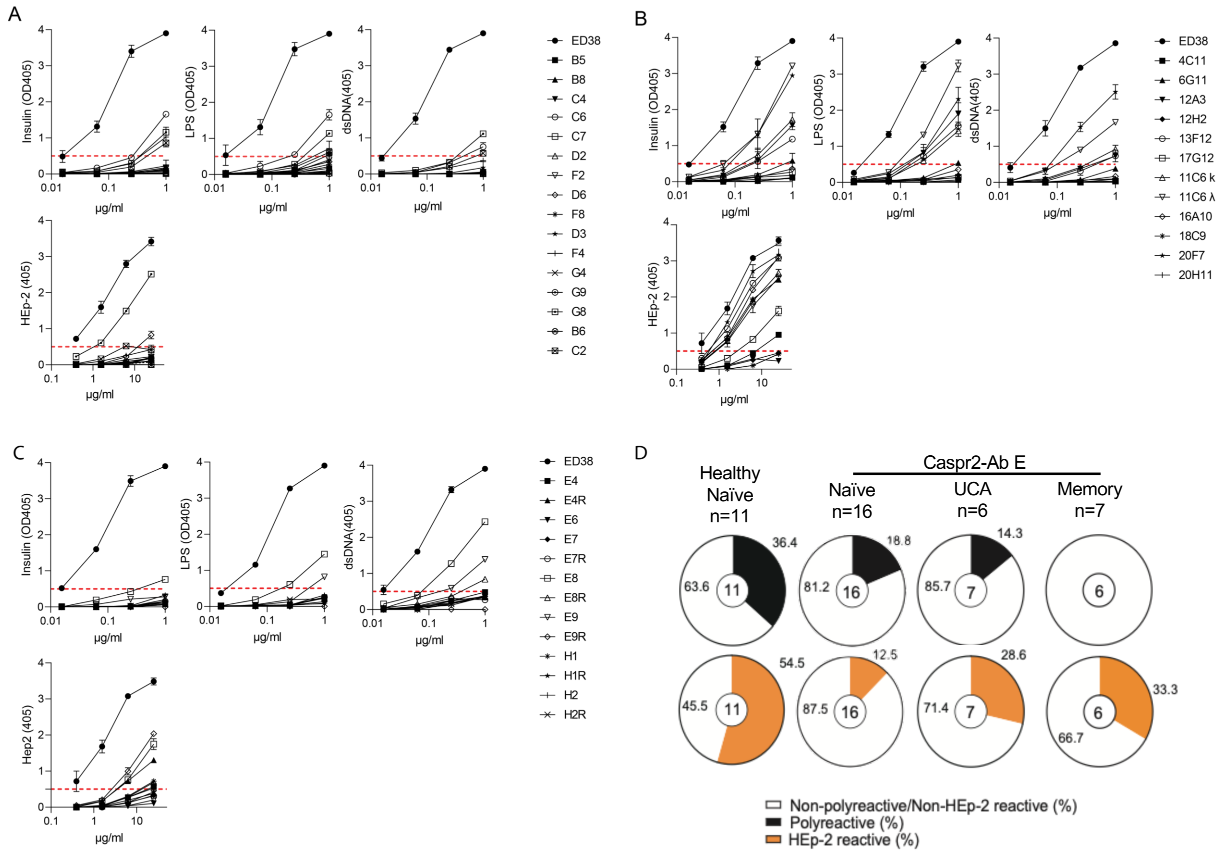


**Figure S4. Insulin/LPS/dsDNA and HEp-2 cell lysate binding by CASPR2 mAbs**
ELISA quantification of mAb binding to insulin/LPS/dsDNA (top) and Hep-2 cell lysate (bottom) by naïve B cell mAbs from CASPR2-Ab E patients (A), healthy control mAbs (B), and memory B cell mAbs from CASPR2-Ab E patients (C). Each data point represents the mean value of two independent experiments and the error bars represent SDs. Dotted horizontal red lines mark the positive reactivity cut-off at OD405 0.5. ED38, a monoclonal antibody cloned from a VpreB + L + peripheral B cell, was a positive control. (D) Pie charts display the overserved frequency distributions for insulin/LPS/dsDNA reactivity (black, left), or Hep-2 reactivity (right, orange). The number in the center of the pie chart represents the total number of unique mAbs tested.

| ID | CASPR2 serostatus (IgG) | CASPR2 serostatus (IgM) | Sex | Age (years) | Disease duration (years) | Clinical Syndrome | Treatment at time of sample collection | Co-morbidities | Total wells NBC (Bulk) | Total wells MBC (Bulk) | Total wells NBC (single-cell) | Total wells MBC (single-cell) |
|---|---|---|---|---|---|---|---|---|---|---|---|---|
| P1 | Positive | Negative | M | 63 | 2 | Cog, seizures | Nil | Nil | 9 | 11 | 2112 | 2208 |
| P2 | Positive | Negative | M | 75 | 7 | Cog, seizures, pain | Nil | BPH, T2DM | 2 | 6 | 2400 | 1598 |
| P3 | Positive | Negative | M | 69 | 1 | Cog, Psy, Pain, PNH | Nil | BPH, HTN, AF | 10 | 1 | - | - |
| P4 | Positive | Negative | M | 75 | 1 | Cog, Psy, seizures | Corticosteroids | BPH, Gout, HTN | 6 | 5 | - | - |
| P5 | Positive | Negative | M | 71 | 0.75 | Cog, Psy, seizures | Corticosteroids | T2DM | 2 | 1 | - | - |
| P6 | Positive | Negative | M | 81 | 7 | Cog, seizures, ANS | Methotrexate | BPH, HTN, AF | 2 | 1 | - | - |
| H1 | Negative | Negative | N/A | N/A | N/A | N/A | N/A | N/A | 25 | 1 | - | - |
| H2 | Negative | Negative | N/A | N/A | N/A | N/A | N/A | N/A | 23 | 4 | - | - |
| H3 | Negative | Negative | N/A | N/A | N/A | N/A | N/A | N/A | 56 | 2 | - | - |
| H4 | Negative | Negative | N/A | N/A | N/A | N/A | N/A | N/A | 30 | 3 | - | - |
| H5 | Negative | Negative | M | 46 | N/A | N/A | N/A | N/A | - | - | 960 | 960 |
| H6 | Negative | Negative | F | 33 | N/A | N/A | N/A | N/A | - | - | 960 | 960 |

Table S1: Patient Demographics

Tabular description of CASPR2-Ab E patients (labeled P1-P6) and healthy controls (labeled H1-H6). M = male, F = female, Cog = cogntive impairment, Psy = psychiatric features, PNH = peripheral nerve hyperexcitability, ANS = autonomic nervous system dysfunction, BPH = benign prostatic hypertrophy, T2DM = Type 2 diabetes mellitus, HTN = hypertension, AF = atrial fibrillation, NBC = naive b cell, MBC = memory b cell, N/A = not applicable, - = not performed

| mAb | Donor | Subset | IGHV | IGHJ | HCDR3 | H DNA Mutations | IG K/L V | IG K/L L | L DNA Mutations |
|---|---|---|---|---|---|---|---|---|---|
| 11C6 | H5 | NBC | IGHV1-18 | IGHJ1*01 | CARDYDFWSGYYTAREYFQHW | 0 | IGKV3-15 | IGKJ4*01 | 0 |
| 16A10 | H5 | NBC | IGHV1-46 | IGHJ6*02 | CAREYGDYVGYYYYGMDVW | 2 | IGLV3-1 | IGLJ2*01 | 0 |
| 18C9 | H5 | NBC | IGHV4-59 | IGHJ4*02 | CARLSYYYGSGNFDYW | 0 | IGLV3-21 | IGLJ1*01 | 0 |
| 20F7 | H5 | NBC | IGHV3-9 | IGHJ6*02 | CAKEGGSYRPAYYYYYYGMDVW | 0 | IGLV3-19 | IGLJ2*01 | 2 |
| 20H11 | H5 | NBC | IGHV3-9 | IGHJ4*02 | CAKETSRDGYNPFDYW | 0 | IGKV1-5 | IGKJ1*01 | 0 |
| 12A3 | H6 | NBC | IGHV3-43D | IGHJ4*02 | CAKETFLPLLGAVAGTSYFDYW | 0 | IGLV3-21 | IGLJ2*01 | 0 |
| 12H2 | H6 | NBC | IGHV3-9 | IGHJ4*02 | CAKEDDYGSLDYW | 0 | IGKV1-17 | IGKJ4*01 | 0 |
| 13F12 | H6 | NBC | IGHV4-31 | IGHJ5*02 | CARDLSGYYYDSSGYYSPIHNWFDPW | 0 | IGKV1-39 | IGKJ1*01 | 0 |
| 17G12 | H6 | NBC | IGHV3-9 | IGHJ6*02 | CAKDSLPGDPYYYYYGMDVW | 0 | IGKV3-20 | IGKJ2*01 | 1 |
| 4C11 | H6 | NBC | IGHV3-7 | IGHJ4*02 | CARFGWAVYGDGRGGFDYW | 0 | IGLV3-25 | IGLJ2*01 | 0 |
| 6G11 | H6 | NBC | IGHV3-33 | IGHJ6*02 | CARDLKPGHQAQAYYYYGMDVW | 0 | IGLV3-19 | IGLJ2*01 | 0 |
| E04 | P1 | MBC | IGHV3-23 | IGHJ4*02 | CAKEGQQQLGFDYW | 10 | IGLV3-1 | IGLJ2*01 | 13 |
| E06 | P1 | MBC | IGHV2-5 | IGHJ5*02 | CAHRPEYSSSWHSWFDPW | 13 | IGKV3-11 | IGKJ2*01 | 15 |
| E07 | P1 | MBC | IGHV3-30 | IGHJ6*02 | CAKEEHGGNKYSYGMDVW | 10 | IGLV1-47 | IGLJ1*01 | 13 |
| E08 | P1 | MBC | IGHV3-23 | IGHJ6*02 | CAKEGFAVGPYSYQYSTMDVW | 21 | IGKV1-39 | IGKJ2*01 | 15 |
| E09 | P1 | MBC | IGHV4-34 | IGHJ4*02 | CARGAYDYVWGNYRYTGVLDYW | 5 | IGKV1-5 | IGKJ2*01 | 4 |
| F02 | P1 | NBC | IGHV3-53 | IGHJ6*02 | CARDEGIQGGYYYYGMDVW | 0 | IGKV1-39 | IGKJ3*01 | 0 |
| F04 | P1 | NBC | IGHV4-31 | IGHJ4*02 | CARFTYYDILTGYSPGFDYW | 0 | IGKV3-20 | IGKJ1*01 | 0 |
| F08 | P1 | NBC | IGHV3-33 | IGHJ4*02 | CAREGRGYSGYGSAGFDYW | 0 | IGLV3-25 | IGLJ3*02 | 6 |
| G04 | P1 | NBC | IGHV1-8 | IGHJ1*01 | CARADFWSGYYSHPW | 0 | IGLV3-1 | IGLJ1*01 | 4 |
| G08 | P1 | NBC | IGHV4-34 | IGHJ6*02 | CASAQGYYDFWSGYYPYYYYGMDVW | 0 | IGLV1-47 | IGLJ1*01 | 2 |
| G09 | P1 | NBC | IGHV3-7 | IGHJ6*02 | CARDGYYDIPYGMDVW | 0 | IGLV3-1 | IGLJ1*01 | 4 |
| H01 | P2 | MBC | IGHV3-23 | IGHJ3*01 | CAKHKDDYDSGSPPIALDVW | 23 | IGLV1-47 | IGLJ1*01 | 18 |
| H02 | P2 | MBC | IGHV3-21 | IGHJ3*01 | CAREYDYADDFLQAAFDVW | 37 | IGLV3-19 | IGLJ3*02 | 29 |
| H03 | P2 | MBC | IGHV3-23 | IGHJ3*01 | CAKHKDDYDSGSPPIALDVW | 23 | IGLV1-47 | IGLJ1*01 | 18 |
| H04 | P2 | MBC | IGHV3-21 | IGHJ3*01 | CAREYDYADDFLQAAFDVW | 37 | IGLV3-19 | IGLJ3*02 | 29 |
| H05 | P2 | MBC | IGHV3-23 | IGHJ3*01 | CAKHKDDYDSGSPPIALDVW | 23 | IGLV1-47 | IGLJ1*01 | 18 |
| B05 | P2 | NBC | IGHV4-61 | IGHJ2*01 | CARDIGYYDSSGYYWYFDLW | 0 | IGLV1-47 | IGLJ3*02 | 2 |
| B06 | P2 | NBC | IGHV3-30 | IGHJ5*02 | CAKDIQSGSGPW | 1 | IGLV3-1*01 | IGLJ1*01 | 0 |
| B08 | P2 | NBC | IGHV3-33 | IGHJ4*02 | CARDRCSGGSCYLLGYFDYW | 0 | IGKV3-11 | IGKJ3*01 | 1 |
| C02 | P2 | NBC | IGHV4-61 | IGHJ6*02 | CAREGYYGSGSSPVYYYYGMDVW | 0 | IGKV1-8 | IGKJ1*01 | 0 |
| C04 | P2 | NBC | IGHV4-31 | IGHJ5*02 | CARSYYYDSSGYYYIWFDPW | 0 | IGLV1-40 | IGLJ3*02 | 3 |
| C06 | P2 | NBC | IGHV3-21 | IGHJ6*02 | CARDRLGYCSGGSCYPLYYYYYGMDVW | 0 | IGKV1-39 | IGKJ1*01 | 1 |
| C07 | P2 | NBC | IGHV3-48 | IGHJ6*02 | CARDGAAAGTNYYYGMDVW | 0 | IGLV3-1 | IGLJ1*01 | 4 |
| D02 | P2 | NBC | IGHV3-7 | IGHJ4*02 | CARVVGATNDIDYW | 0 | IGKV1-39 | IGKJ2*01 | 1 |
| D03 | P2 | NBC | IGHV3-23 | IGHJ6*02 | CAKEGQYYDFWSGYSAVDYYYYGMDVW | 0 | IGLV3-1 | IGLJ1*01 | 5 |
| D06 | P2 | NBC | IGHV3-9 | IGHJ3*02 | CAKDTHSLQNAFDIW | 0 | IGKV1-39 | IGKJ1*01 | 5 |

Table S2: Recovered CASPR2 mAbs
Tabular description of recovered mAbs , with structural information obtained by IMGT High Vquest. NBC = naïve B cell, MBC = memory B cell

| Antibody | Ka ($M^{-1}s^{-1}$) | Kd ($s^{-1}$) | $K_D$ (M) | t 1/2 (s) |
|---|---|---|---|---|
| E09 | 3.04E+05 | 4.64E-05 | 1.53E-10 | 1.49E+04 |
| E09-UCA | 5.09E+04 | 1.70E-04 | 3.33E-09 | 4.08E+03 |
| E08 | 3.70E+08 | 1.11E+01 | 3.00E-08 | 6.25E-02 |
| E08-UCA | 6.38E+04 | 8.92E-02 | 1.40E-06 | 7.77E+00 |
| E07 | 3.14E+03 | 1.54E-04 | 4.90E-08 | 4.51E+03 |
| E07-UCA | 2.60E+03 | 3.99E-03 | 1.54E-06 | 1.74E+02 |
| H02 | 4.79E+05 | 2.62E-01 | 5.47E-07 | 2.64E+00 |
| H02-UCA | ND | ND | ND | ND |
| H01 | 1.63E+07 | 1.37E+01 | 8.37E-07 | 5.07E-02 |
| H01-UCA | ND | ND | ND | ND |
| E04 | 3.21E+04 | 1.72E-01 | 5.37E-06 | 4.03E+00 |
| E04-UCA | ND | ND | ND | ND |
| E06 | 1.08E+03 | 2.89E-01 | 2.67E-04 | 2.40E+00 |

Table S3: Surface plasma resonance binding quantifications of memory B cell derived mAbs and their corresponding UCAs. Binding not detected = ND

| Name | Chain | Primer Sequences (5’ to 3’) | PCR Reaction |
|---|---|---|---|
| CHG1 | IgG H | GTTGTCCACCTTGGTGTTGCTGG | 1 |
| VHL1 | IgG,M H | CCATGGACTGGACCTGGAG | 1 |
| VHL1 mod | IgG M H | CCATGGACTGSACCTGGAG | 1 |
| VHL2 | IgG,M H | ATGGACATACTTTGCTCCAC | 1 |
| VHL2 mod | IgG M H | ATGGACATACTTTGYTCCAC | 1 |
| VHL3 | IgG,M H | CCATGGAGTTKGGGCTGAGCTGG | 1 |
| VHL4 | IgG,M H | ATGAAACACCTGTGGTTCTT | 1 |
| VHL4 mod | IgG M H | ATGAAACAYCTGTGGTTCTT | 1 |
| VHL5 | IgG,M H | ATGGGGTCAACCGCCATCCT | 1 |
| CHm1 | IgG M | GAAGCCAGCACCTGTGAGG | 1 |
| CK1 | Kappa | ACACTCTCCCCTGTTGAAGCTCTT | 1 |
| VKL1 | Kappa | CTCAGCTCCTGGGGCTCC | 1 |
| VKL1 mod | Kappa | CTCAGCTCCTGGGGCTYC | 1 |
| VKL2 | Kappa | CTGCTCAGCTCCTGGGGC | 1 |
| VKL2 mod | Kappa | CTGCTCAGCTCYTGGGGC | 1 |
| VKL3 | Kappa | GGAARCCCCAGCDCAGC | 1 |
| VKL4 | Kappa | CTCTGTTGCTCTGGATCTCTG | 1 |
| VKL5 | Kappa | GGGGTCCCAGGTTCACCTCCTC | 1 |
| CL1 | Lambda | TGAACATTCTGTAGGGGCCAC | 1 |
| VLL1 | Lambda | CCTCTCCTCCTCACCCTCCTC | 1 |
| VLL1 mod | Lambda | CCTCTCCTCCTSACCCTCCTC | 1 |
| VLL2 | Lambda | CTCCTCACTCAGGGCACAG | 1 |
| VLL2 mod | Lambda | CTCCTCACTCAGGRCACAG | 1 |
| VLL3 | Lambda | ATGGCCTGGAYCCCTCTCCTSCT | 1 |
| VLL3 mod | Lambda | ATGGCCTGGAYCCCTCTCCTBCT | 1 |
| VLL4a | Lambda | CTCCTCCTCTTCCCCCTCCCCCTC | 1 |
| VLL4a mod | Lambda | CTCCTCCTCCACTGCACAG | 1 |
| VLL4b | Lambda | TGGCCTGGGTCTCCTTCTACCTAC | 1 |
| VLL569 | Lambda | ATGGCCTGGRCTCCTCTCCTYCTC | 1 |
| VLL710 | Lambda | ATGGCCTGGRCTCCTCTCCTYCTG | 1 |
| VLL8 | Lambda | ATGGCCTGGATGATGCTTCTCCTC | 1 |
| CPEC VH1/5/7 | Heavy | CTTTTTCTAGTAGCAACTGCAACCGGTGTACATTCCGAGGTGCAGCTGGTGCAG | 2 |
| CPEC VH2 | Heavy | CTTTTTCTAGTAGCAACTGCAACCGGTGTACATTCCCAGGTCACCTTGAAGGAG | 2 |
| CPEC VH3 | Heavy | CTTTTTCTAGTAGCAACTGCAACCGGTGTACATTCTGAGGTGCAGCTGGTGGAG | 2 |
| CPEC VH4 | Heavy | CTTTTTCTAGTAGCAACTGCAACCGGTGTACATTCCCAGGTGCAGCTGCAGGAG | 2 |
| CPEC VH3-23 | Heavy | CTTTTTCTAGTAGCAACTGCAACCGGTGTACATTCTGAGGTGCAGCTGTTGGAG | 2 |
| CPEC VH4-34 | Heavy | CTTTTTCTAGTAGCAACTGCAACCGGTGTACATTCCCAGGTGCAGCTACAGCAGTG | 2 |
| CPEC VH1–18/69 (silent | Heavy | CTTTTTCTAGTAGCAACTGCAACCGGTGTACATTCCCAGGTTCAGCTGGTGCAG | 2 |
| CPEC VH1-45/1-58 | Heavy | CTTTTTCTAGTAGCAACTGCAACCGGTGTACATTCCCAGATGCAGCTGGTGCAG | 2 |
| CPEC VH1–24 | Heavy | CTTTTTCTAGTAGCAACTGCAACCGGTGTACATTCCCAGGTCCAGCTGGTACAG | 2 |
| CPEC VH3–9/30/33 | Heavy | CTTTTTCTAGTAGCAACTGCAACCGGTGTACATTCTGAAGTGCAGCTGGTGGAG | 2 |
| CPEC VH6–1 | Heavy | CTTTTTCTAGTAGCAACTGCAACCGGTGTACATTCCCAGGTACAGCTGCAGCAG | 2 |
| CPEC VH4–39 | Heavy | CTTTTTCTAGTAGCAACTGCAACCGGTGTACATTCCCAGCTGCAGCTGCAGGAG | 2 |
| CPEC VH3–33/11/30 | Heavy | CTTTTTCTAGTAGCAACTGCAACCGGTGTACATTCTCAGGTGCAGCTGGTGGAG | 2 |
| CPEC JH1/2/4/5 | Heavy | GATGGGCCCTTGGTCGACGCTGAGGAGACGGTGACCAG | 2 |
| CPEC JH3 | Heavy | GATGGGCCCTTGGTCGACGCTGAAGAGACGGTGACCATTG | 2 |
| CPEC JH6 | Heavy | GATGGGCCCTTGGTCGACGCTGAGGAGACGGTGACCGTG | 2 |
| CPEC VK1 | Kappa | CTTTTTCTAGTAGCAACTGCAACCGGTGTACATTCTGACATCCAGATGACCCAGTC | 2 |
| CPEC VK1-9/1-13 | Kappa | CTTTTTCTAGTAGCAACTGCAACCGGTGTACATTCAGACATCCAGTTGACCCAGTCT | 2 |
| CPEC VK1D-43/1-8 | Kappa | CTTTTTCTAGTAGCAACTGCAACCGGTGTACATTGTGCCATCCGGATGACCCAGTC | 2 |
| CPEC VK2 | Kappa | CTTTTTCTAGTAGCAACTGCAACCGGTGTACATGGGGATATTGTGATGACCCAGAC | 2 |
| CPEC VK2-28/2-30 | Kappa | CTTTTTCTAGTAGCAACTGCAACCGGTGTACATGGGGATATTGTGATGACTCAGTC | 2 |
| CPEC VK3-11/3D-11 | Kappa | CTTTTTCTAGTAGCAACTGCAACCGGTGTACATTCAGAAATTGTGTTGACACAGTC | 2 |
| CPEC VK3-15/3D-15 | Kappa | CTTTTTCTAGTAGCAACTGCAACCGGTGTACATTCAGAAATAGTGATGACGCAGTC | 2 |
| CPEC VK3-20/3D-20 | Kappa | CTTTTTCTAGTAGCAACTGCAACCGGTGTACATTCAGAAATTGTGTTGACGCAGTCT | 2 |
| CPEC VK4-1 | Kappa | CTTTTTCTAGTAGCAACTGCAACCGGTGTACATTCGGACATCGTGATGACCCAGTC | 2 |
| CPEC JK1/2/4 | Kappa | ATGGTGCAGCCACCGTACGTTTGATYTCCACCTTGGTC | 2 |
| CPEC JK2 | Kappa | ATGGTGCAGCCACCGTACGTTTGATCTCCAGCTTGGTC | 2 |
| CPEC JK3 | Kappa | ATGGTGCAGCCACCGTACGTTTGATATCCACTTTGGTC | 2 |
| CPEC JK5 | Kappa | ATGGTGCAGCCACCGTACGTTTAATCTCCAGTCGTGTC | 2 |
| CPEC JK150/3 | Kappa | ATGGTGCAGCCACCGTACGTCTGATTTCCACCTTGGTC | 2 |
| CPEC VL1 | Lambda | CTTTTTCTAGTAGCAACTGCAACCGGTTCCTGGGCCCAGTCTGTGCTGACKCAG | 2 |
| CPEC VL2 | Lambda | CTTTTTCTAGTAGCAACTGCAACCGGTTCCTGGGCCCAGTCTGCCCTGACTCAG | 2 |
| CPEC VL3 | Lambda | CTTTTTCTAGTAGCAACTGCAACCGGTTCTGTGACCTCCTATGAGCTGACWCAG | 2 |
| CPEC VL4/5 | Lambda | CTTTTTCTAGTAGCAACTGCAACCGGTTCTCTCTCSCAGCYTGTGCTGACTCA | 2 |
| CPEC VL6 | Lambda | CTTTTTCTAGTAGCAACTGCAACCGGTTCTTGGGCCAATTTTATGCTGACTCAG | 2 |
| CPEC VL7/8 | Lambda | CTTTTTCTAGTAGCAACTGCAACCGGTTCCAATTCYCAGRCTGTGGTGACYCAG | 2 |
| CPEC CL reverse | Lambda | GGCTTGAAGCTCCTCACTCGAGGGYGGGAACAGAGTG | 2 |
| 3’CL Reverse | Lambda | CACCAGTGTGGCCTTGTTGGCTTG | 2 |
| panVK Forw. | Kappa | ATGACCCAGWCTCCABYCWCCCTG | 2 |
| 3’CK494-516 R. | Kappa | GTGCTGTCCTTGCTGTCCTGCT | 2 |
| 3’ CgCH Reverse Tiller | Heavy | GGAAGGTGTGCACGCCGCTGGTC | 2 |

Table S4: Primer name, target, sequence and PCR reaction from lysed B cells

| Target | Clone or laboratory number; Manufacturer | Assays | Concentration or Dilution |
|---|---|---|---|
| CD3 | UCHT1; BioLegend | FACS | 1:80 |
| CD14 | HCD14; BioLegend | FACS | 1:80 |
| CD19 | SJ25C1; BD Biosciences | FACS | 1:32 |
| CD27 | O323; BioLegend | FACS | 1:16 |
| IgD | IA6-2; BD Biosciences | FACS | 1:40 |
| CD38 | HB7; BD Biosciences | FACS | 1:40 |
| DAPI | 564907; BD Pharmingen | FACS; Cell based assays | FACS: 0.5 ug/ml, CBA: 1:5000 |
| Human IgG | A11014; Thermofisher | Cell based assays | 1:750 |
| Human IgM | A21216; Thermofisher | Cell based assays | 1:500 |
| Human IgG | SAB3701279; Sigma | Brain sections | 1:500 |
| Caspr2 | Ab33994; Abcam | Cortical rat neurons | 1:200 |
| MAP2 | Ab5392; Abcam | Cortical rat neurons | 1:5000 |
| PSD-95 | 3450; Cell Signalling Technologies | Cortical rat neurons | 1:750 |
| pan-GluA | 182411; Synaptic Systems | Cortical rat neurons | 1:100 |
| Chicken IgY | 103-155-155; Jackson ImmunoResearch | Cortical rat neurons | 1:200 |
| Mouse IgG | A11001; Thermofisher | Cortical rat neurons | 1:500 |
| Rabbit IgG | A11036; Thermofisher | Cortical rat neurons | 1:500 |
| Human IgG | A-11013ThermoFisher | Dorsal root ganglion rat | 5 µg/mL |
| Beta tubulin III | Ab179511; Abcam | Dorsal root ganglion rat | 2 µg/mL |
| Human IgG | 31125; ThermoFisher | Hippocampal rat neurons | 2 and 3 µg/mL |
| Goat IgG | A11055; ThermoFisher | Hippocampal rat neurons | 1:750 |
| MAP2 | ab5392; Abcam | Hippocampal rat neurons | (1:3000) |
| Chicken IgY | A11041; ThermoFisher | Hippocampal rat neurons | 3 µg/mL |
| Human IgG (Fc) | SA5-10134; ThermoFisher | Sensory-neuron IPSCs | 1:500 |
| Goat IgG | A11055, ThermoFisher | Sensory-neuron IPSCs | 1:500 |
| Beta tubulin III | T2200; Sigma | Sensory-neuron IPSCs | 1:2000 |
| Rabbit IgG | P-10994; ThermoFisher | Sensory-neuron IPSCs | 1:1000 |

Table S5: List of commercial antibodies
